# Septins and K63 chains form separate bacterial microdomains during autophagy of entrapped *Shigella*

**DOI:** 10.1101/2022.11.14.516380

**Authors:** Damián Lobato-Márquez, José Javier Conesa, Ana Teresa López-Jiménez, Michael E. Divine, Jonathan N. Pruneda, Serge Mostowy

**Affiliations:** Department of Infection Biology, London School of Hygiene and Tropical Medicine, Keppel Street, London WC1E 7HT, United Kingdom; Department of Structure of Macromolecules, Centro Nacional de Biotecnología, CSIC, 28049 Madrid, Spain; MISTRAL beamline, ALBA Synchrotron Light Source, Cerdanyola del Vallès, 08290 Barcelona, Spain; Department of Molecular Microbiology & Immunology, Oregon Health & Science University, Portland, OR 97239, USA

**Keywords:** autophagy, cytoskeleton, cryo-SXT, septins, *Shigella*, ubiquitin

## Abstract

During host cell invasion, *Shigella* escapes to the cytosol and polymerizes actin for cell-to-cell spread. To restrict cell-to-cell spread, host cells employ cell-autonomous immune responses including antibacterial autophagy and septin cage entrapment. How septins interact with autophagy to target *Shigella* to destruction is poorly understood. Here, we employed a correlative light and cryo-soft X-ray tomography (cryo-SXT) pipeline to study *Shigella* septin cage entrapment in its near native state. Quantitative cryo-SXT showed that *Shigella* fragments mitochondria and enabled visualization of X-ray dense structures (∼30 nm resolution) surrounding *Shigella* entrapped in septin cages. Using Airyscan confocal microscopy, we observed Lysine 63 (K63)-linked ubiquitin chains decorating septin caged entrapped *Shigella*. Remarkably, septins and K63 chains form separate bacterial microdomains, indicating they are recruited separately during antibacterial autophagy. Cryo-SXT and live cell imaging revealed an interaction between septins and LC3B-positive membranes during autophagy of *Shigella*. Together, these findings demonstrate how septin caged *Shigella* are targeted to autophagy and provide fundamental insights into autophagy-cytoskeleton interactions.

## Introduction

Septins are an evolutionarily conserved family of GTP-binding proteins that interact with cellular membranes to form nonpolar filaments and higher order ring-like structures (Spiliotis and McMurray, 2020; Woods and Gladfelter, 2020). Septin interactions with the plasma membrane underpin a variety of eukaryotic cell hallmarks including the cytokinetic furrow, cellular protrusions (e.g., cilium and dendritic spines) and the phagocytic cup surrounding invasive bacterial pathogens (Lobato-Marquez and Mostowy, 2016; Mostowy and Cossart, 2012). In the cytosol, septins can entrap *S. flexneri* in cage-like structures to restrict actin based motility (Mostowy et al., 2010), and mitochondria can promote this process (Sirianni et al., 2016). To counteract septin cage entrapment, actin-polymerizing *S. flexneri* can fragment mitochondria (Sirianni et al., 2016). Our previous work has shown that septins recognize micron-scale curvature presented by bacterial membranes enriched in cardiolipin (Krokowski et al., 2018). More recently, we developed an *in vitro* reconstitution assay using purified proteins to dissect mechanisms underlying septin recognition of growing bacterial cells (Lobato-Márquez et al., 2021). Despite recent insights, the role of septin interactions with host cell membrane during *S. flexneri* cage entrapment is poorly understood.

In addition to restriction of actin-based motility, septin cages target entrapped bacteria to destruction by autophagy (Krokowski et al., 2018; Mostowy et al., 2010; Sirianni et al., 2016). Autophagy, an evolutionarily conserved degradative process that breaks down cytosolic material inside double-membrane vesicles (autophagosomes) by fusion with lysosomes, has key roles in maintaining cellular homeostasis, recycling damaged organelles and providing nutrients during starvation (Bento et al., 2016; Dikic and Elazar, 2018). Selective autophagy is an important host defense mechanism that recognizes intracellular bacterial pathogens for degradation (also called xenophagy) (Grumati and Dikic, 2018). The best described mechanism underlying selective autophagy is via ubiquitination of host or bacterial proteins (Randow et al., 2013). Ubiquitination is a highly versatile posttranslational modification regulating a wide variety of fundamental cellular processes. Ubiquitin is a small protein (76 amino acids) which contains 7 lysine residues and an N-terminal methionine residue that can be attached to another ubiquitin monomer (Hershko and Ciechanover, 1998); in this way, proteins can be modified with different polyubiquitin lengths and linkages that direct distinct signaling outcomes (Yau and Rape, 2016). In the case of xenophagy, components of *Salmonella enterica* serovar Typhimurium and *Mycobacterium tuberculosis*-containing phagosomes are targeted with lysine 63 (K63)-linked ubiquitin chains (Fiskin et al., 2016; Manzanillo et al., 2013; van Wijk et al., 2012). Autophagy adaptor proteins (such as p62,

NDP52, OPTN) can bind ubiquitin and recruit microtubule-associated protein light chain 3B (LC3B), an important component of the canonical autophagy machinery (Huang and Brumell, 2014; Klionsky et al., 2021; Thurston et al., 2009; Wild et al., 2011; Zheng et al., 2009). In the case of HeLa cell infection, *S. flexneri* septin cages are known to co-localize with ubiquitin (Mostowy et al., 2010; Mostowy et al., 2011). New work performed *in vitro* using purified proteins has shown that septins can directly bind the outer membrane of growing *S. flexneri* cells (Lobato-Márquez et al., 2021). However, whether septins or bacterial membrane components are ubiquitinated, and the type of ubiquitin linkage employed by the host cell during cage entrapment was unknown. While the bacterial septin cage is considered a paradigm for the investigation of cytoskeleton-autophagy interactions (Mostowy and Shenoy, 2015; Van Ngo and Mostowy, 2019; Welch and Way, 2013), we still lack fundamental understanding of how septin-caged entrapped bacterial cells are targeted to autophagy.

Recent advances in microscopy have revolutionized cellular microbiology and the study of host-pathogen interactions (López-Jiménez and Mostowy, 2021). Cryo-soft X-ray tomography (cryo-SXT) is a technique that permits imaging of unstained and cryopreserved biological samples (in their near native state) as thick as 10 µm, therefore overcoming most EM limitations (Carrascosa et al., 2009; Conesa et al., 2016; Harkiolaki et al., 2018; Schneider et al., 2010). In addition, cryo-SXT possess a resolving power in the nanometer scale (as high as 30 nm) and high contrast for biological samples, making cryo-SXT an ideal technique to study the precise localization and organization of membrane-based organelles and pathogens within the host cell cytosol (Carrascosa et al., 2009; Conesa et al., 2020; Cruz-Adalia et al., 2014; Groen et al., 2019; Kounatidis et al., 2020). In this report, we use correlative light and cryo-SXT, Airyscan confocal microscopy and live cell imaging to study septin-autophagy interactions during *S. flexneri* cage entrapment.

## Results

### Visualization of mitochondrial morphology during *S. flexneri* infection by correlative fluorescence and cryo-SXT

We applied correlative light and cryo-SXT to *S. flexneri* infected HeLa cells (Fig. S1 and S2). Given that *S. flexneri* is reported to fragment mitochondria during host cell invasion (Carneiro et al., 2009; Sirianni et al., 2016), we first studied mitochondrial morphology during bacterial infection. We stained HeLa cells with Mitotracker Red and infected them (or not) with *S. flexneri* for 3 h. In the absence of infection, Airyscan confocal microscopy showed that HeLa cells possess elongated mitochondria (Fig. S3) and cryo-SXT demonstrated elongated mitochondria forming an intricate network (Fig. 1A left panels). By contrast, infection with *S. flexneri* caused mitochondrial fragmentation (Fig. 1A right panels and S3); in this case mitochondria are dramatically smaller and less interconnected as compared to uninfected cells. We quantified the length of mitochondria using both fluorescence microscopy and cryo-SXT. Consistent with our previous report that *S. flexneri* fragments mitochondria during host cell invasion (Sirianni *et al*., EMBO Rep 2016), *S. flexneri*-infected HeLa cells showed significantly shorter mitochondria as compared to uninfected HeLa cells (Fig. 1B and 1C). Of note, in the absence of fluorescence markers labelling cytosolic bacteria, the identification of *S. flexneri* was obvious by cryo-SXT (Fig. 1 and Fig. S2). Together, these data demonstrate that *S. flexneri* fragments mitochondria during infection and highlights the potential of cryo-SXT to study other intracellular hallmarks of the *S. flexneri* infection process.

**Figure 1.**
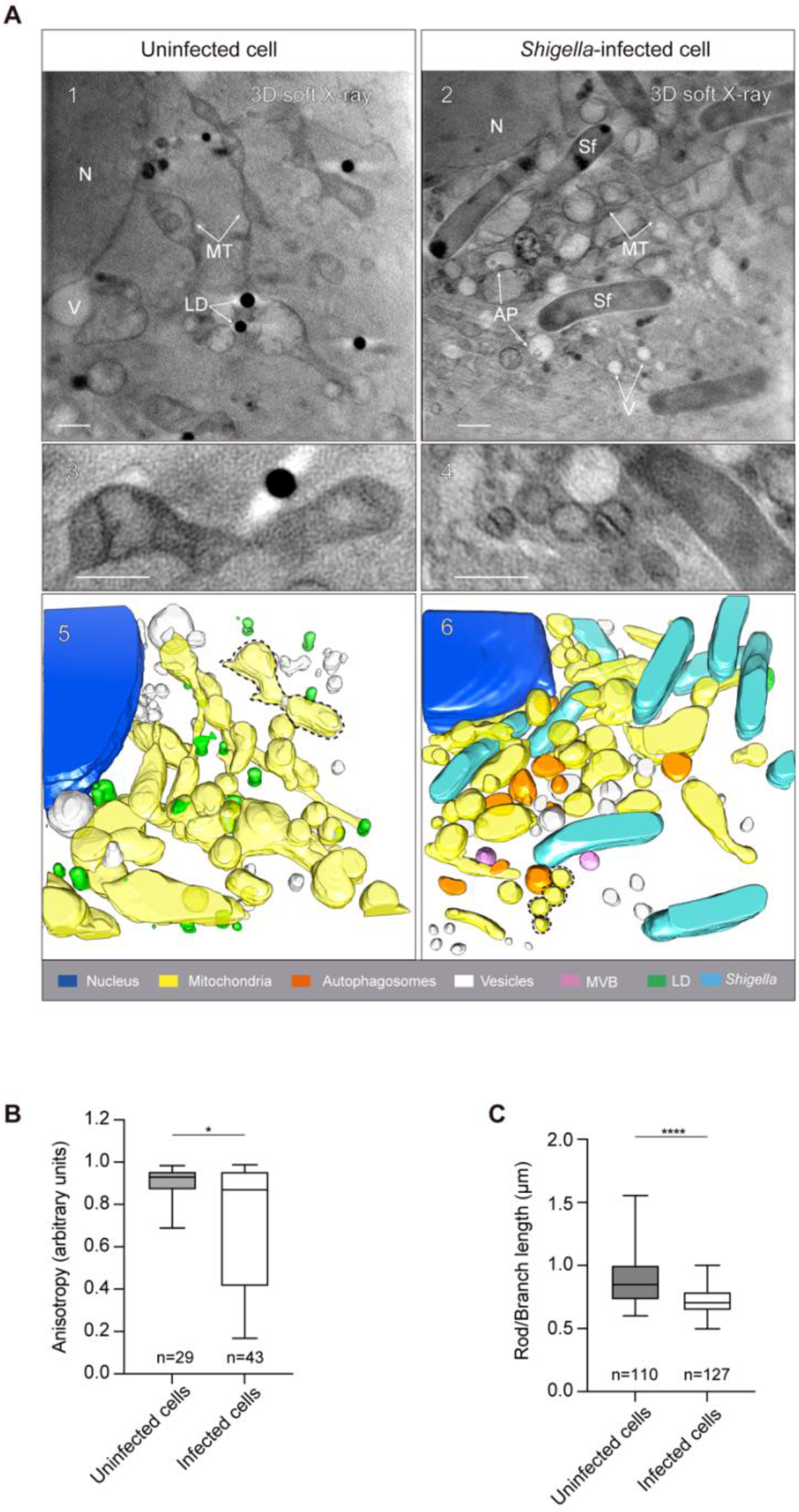
Visualization of mitochondrial dynamics during *Shigella flexneri* infection by correlative fluorescence and cryo-SXT. **(A)** Uninfected (left panels) and *S. flexneri*-infected (right panels) HeLa cells where stained with Mitotracker Red and imaged by correlative light and cryo-SXT. **(1, 2)** Tomographic slices showing elongated mitochondrial network in uninfected cells (left panel), and fragmented mitochondria during *S. flexneri* infection (right panel). Images shown corresponds to slices of 3.9 µm (1) and 4.1 µm thickness. AP, autophagosome; LD, lipid droplet; MT, mitochondria; N, nucleus; V, vesicles. **(3, 4)** Magnification of mitochondria from (1) and (2). **(5, 6)** Volumetric representation of the tomograms in (1) and (2). Scale bars, 1 μm. **(B)** Quantification of mitochondrial elongation using cryo-SXT. Data represent the median ± min/max of anisotropy from n = 29 mitochondria (uninfected cells) from 2 independent tomograms and n = 43 mitochondria (*S. flexneri*-infected cells) from 3 independent tomograms. *, p = 0.038 by Mann-Whitney test. **(C)** Quantification of mitochondrial branch length (a measurement of mitochondrial network elongation) using epifluorescence microscopy and Fiji plugin MiNa. Data represent the median ± min/max of mitochondrial rod/branch length from n = 109 mitochondria (uninfected cells) and n = 127 mitochondria (*S. flexneri*-infected cells) distributed in 2 independent experiments. ****, p < 0.0001 by Mann-Whitney test.

### Use of cryo-SXT to study *S. flexneri* entrapment in septin cages *in situ*

The *S. flexneri* septin cage has been studied for over 10 years using tissue culture cells, zebrafish infection models and a wide variety of fluorescent microscopy techniques (Krokowski et al., 2018; Mostowy et al., 2010; Mostowy et al., 2013; Mostowy et al., 2011; Sirianni et al., 2016). More recently, we imaged bacterial septin cages reconstituted *in vitro* (using purified proteins) at the nanometer scale using cryo-electron tomography, and in this case resolved how septins interact with bacterial membrane (Lobato-Márquez et al., 2021). Despite these efforts, septin cages have not been imaged at high-resolution in their native state during host cell infection. To address this, we employed correlative light and cryo -SXT to visualize septin cage entrapment of *S. flexneri in situ*. HeLa cells producing GFP-SEPT6 (Sirianni et al., 2016) were infected for 3 h with *S. flexneri*, plunge-freezed and introduced into our correlative pipeline (Fig. S1). Bacterial septin cages identified by epifluorescence microscopy were subsequently imaged by X-ray tomography. Strikingly, 92.7% of *S. flexneri* cells entrapped in septin cages could be identified by cryo-SXT as an X-ray dense structure (Fig. 2A, top panel, and Fig. S1). In these images, GFP-SEPT6 fluorescence co-localizes with the X-ray dense structure and when GFP-SEPT6 fluorescence is absent the X-ray dense signal is also absent, strongly suggesting that dark features surrounding bacterial membrane correspond to structures (probably containing host lipids) that are enriched in septins (Fig. 2A top panel, Fig. S4). Consistent with this, the use of correlative light and cryo-SXT revealed that only 11% of bacteria not clearly entrapped in GFP-SEPT6 septin cages show an X-ray dense structure (Fig. 2A bottom panel and 2C).

**Figure 2.**
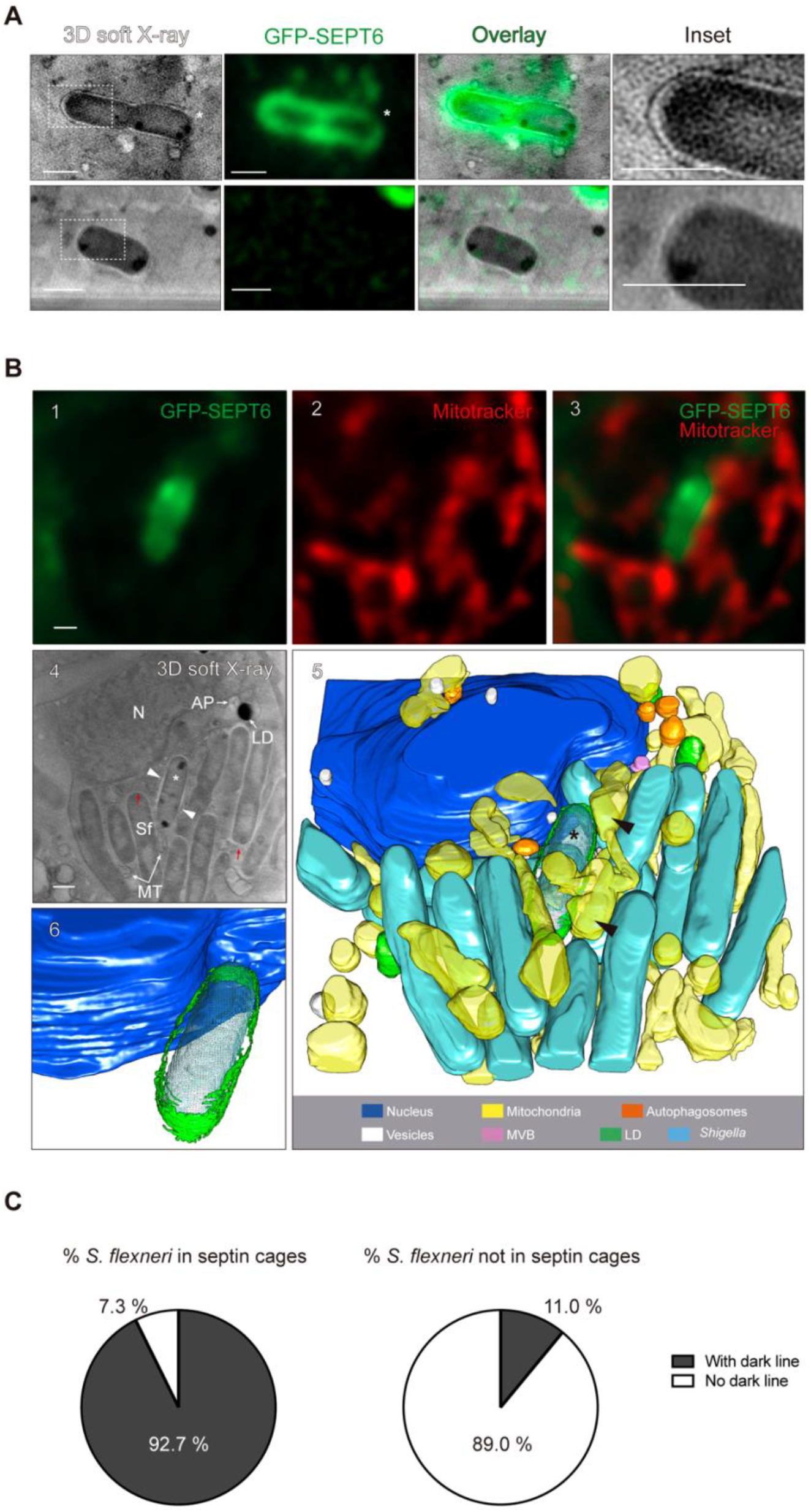
Fluorescent signal of septin cages correlates to increased soft X-ray densities. **(A)** Representative example of a bacterium entrapped in a septin cage imaged by correlative light and cryo-SXT. Soft X-ray dense lines correlate with GFP-SEPT6 fluorescence (top panel inset). Note that where the septin signal is weaker the X-ray signal is also less intense (white asterisk, top panel). Soft X-ray dense lines are not observed around non-caged bacteria (bottom panel). See also Fig. S4. Scale bars, 1 μm. **(B)** Representative cellular environment of a septin caged bacterium. From left to right and top to bottom, (1-3) fluorescence microscopy of septin caged bacterium. (4) Tomographic slice of the same area imaged on (1-3). Image shown corresponds to a slice of 4.6 µm thickness. See also Movie S1. AP, autophagosome; N, nucleus; LD, lipid droplets; MT, mitochondria; Sf, *S. flexneri*; red arrows point to extended bacterial periplasm at the pole of some cells; white arrow heads point to the dense X-ray structure surrounding septin caged *S. flexneri*; *, septin caged bacterium. Note elongated mitochondria surrounding the septin cage entrapped bacterium. (5) Volumetric representation of the tomogram on panel (4). (6) Inset showing the 3D architecture of the *S. flexneri* septin cage, highlighting septins surrounding the entrapped bacterium. Scale bars, 1 μm. See also Movie S1. **(C)** Percentage of *S. flexneri* entrapped or not in septin cages (as defined by fluorescence microscopy) that show a dense X-ray structure surrounding the bacterial cell. Data represent n = 55 (septin caged *S. flexneri* of which 51 show dense X-ray structure) and n = 109 (non-caged *S. flexneri* of which only 12 show dense X-ray structure) bacteria distributed in 53 tomograms.

In contrast to the dense X-ray structures surrounding *S. flexneri* colocalizing with the GFP-SEPT6 fluorescence signal of the septin cage, we occasionally observed a thin X-ray signal at the pole of some bacterial cells (Fig. 2B, 4 red arrows). We hypothesized that this thin X-ray signal, which does not colocalize with GFP-SEPT6 signal, may correspond to the outer membrane of bacteria that exhibit an extended periplasm. To test this, we combined our *in vitro* septin cage reconstitution assay using purified septin complexes (SEPT2–GFP-SEPT6– SEPT7) (Lobato-Márquez et al., 2021) with correlative light and cryo-SXT. As expected, recombinant septin proteins did not provide enough density to be visualized by cryo-SXT (Fig. S4B), reinforcing our conclusion that the X-ray dense structures surrounding bacterial cells are enriched in host cell-derived lipids. In support of our hypothesis that thin X-ray signal present at some bacterial cell poles represents bacterial outer membrane (with an extended periplasm), we could visualize *S. flexneri* outer membrane by cryo-SXT using bacteria grown in broth (Fig. S4B). Consistent with this, previous work has shown that the periplasm of Gram-negative bacteria expands at the bacterial cell poles under stressful conditions (such as bacteria grown in minimal medium or high osmolarity) (Sochacki et al., Biophys J 2011).

Mitochondria promote *S. flexneri* septin cage assembly (Sirianni et al., 2016). In agreement with this, epifluorescence microscopy showed that septin cages are surrounded by elongated mitochondria (Fig. 2B, panels 1-3, Movie S1), and cryo-SXT revealed that those septin-caged bacteria are tightly associated with an elongated mitochondrial network (Fig. 2B, panels 4 and 5, Movie S1). These observations are in sharp contrast with the cellular environment of *S. flexneri* not entrapped by septin cages (Fig. 1), where mitochondria are clearly more fragmented and the mitochondrial network less extensive. Together, cryo-SXT enables the *in situ* visualization of *S. flexneri* septin cages as X-ray-dense structures in close contact with elongated host cell mitochondria.

### Interaction of septins with LC3B-positive membranes during *S. flexneri* entrapment

How septins interact with the autophagy machinery to clear *S. flexneri* is unknown. During visualization of *S. flexneri*-septin cages by correlative light and cryo-SXT we observed septin– autophagosome interactions (Fig. 3A). To confirm that vesicles contacting caged bacterial cells are autophagosomes, we infected HeLa cells stably producing GFP-SEPT6 and transfected with mCherry-LC3B with *S. flexneri* and performed correlative light and cryo-SXT. We observed LC3B-positive vesicles recruited to septin-caged bacteria, supporting the hypothesis that septins and autophagic membranes interact during autophagy of entrapped *S. flexneri* (Fig. 3B, Movie S2). In all cases, *S. flexneri* septin cages are tightly bound to host cell mitochondria (Fig. 3A and 3B). To visualize septin – autophagosome interaction in real time, we performed time-lapse epifluorescence microscopy using HeLa cells producing GFP-SEPT6 and mCherry-LC3B. Consistent with the model that septin cage entrapment and autophagy are interdependent processes (Mostowy et al., 2010; Mostowy et al., 2011), live cell imaging showed a coordinated recruitment of GFP-SEPT6 and mCherry-LC3B to cytosolic *S. flexneri* (Fig. S5, Movie S3 and S4).

**Figure 3.**
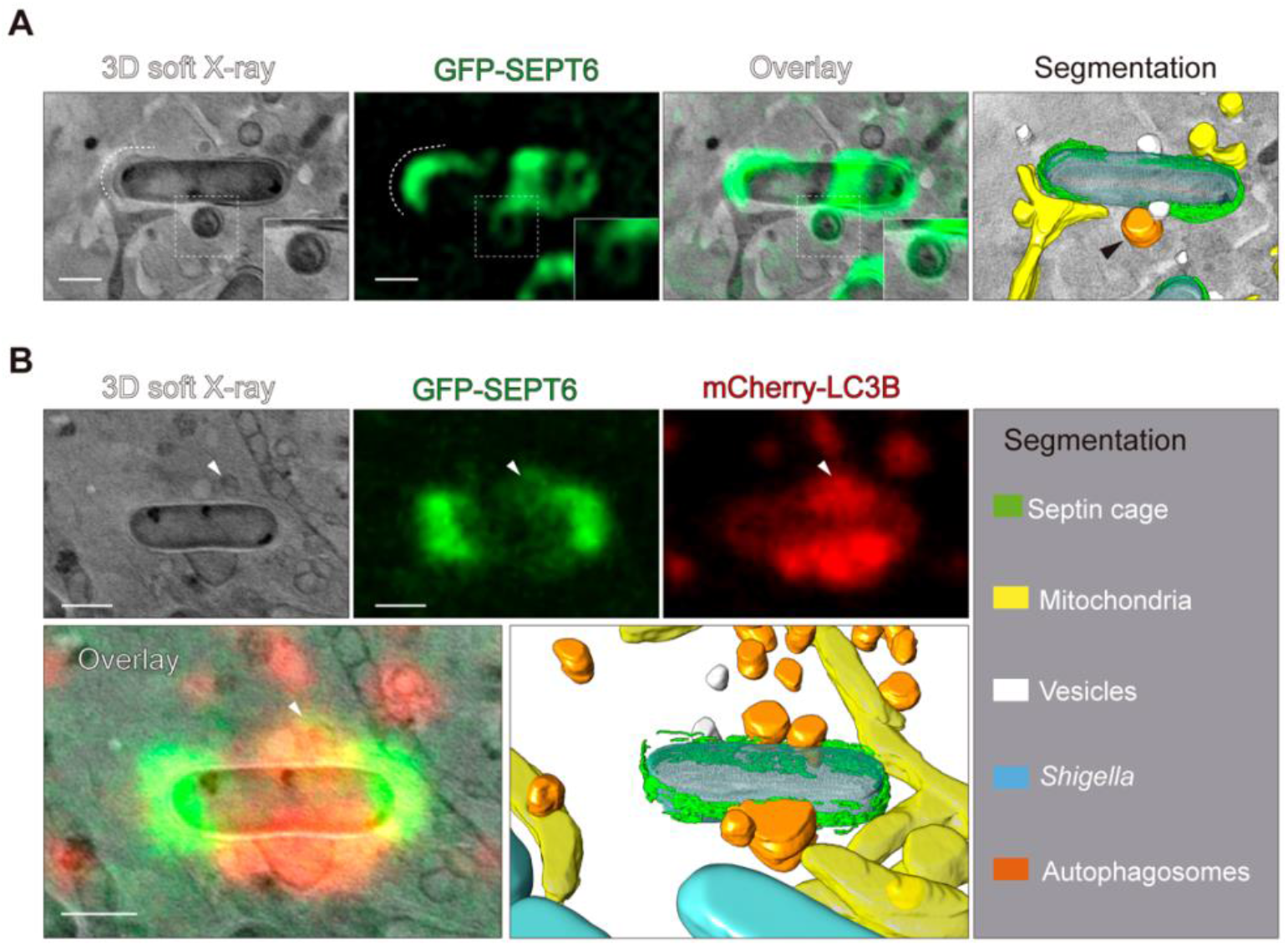
Septins interact with LC3B-decorated vesicles during autophagy of *S. flexneri*. **(A)** Correlative light and cryo-SXT showing septins localize to vesicles interacting with septin-caged *S. flexneri* (black arrow). Scale bar, 1 μm. **(B)** Correlative light and cryo-SXT showing septins localize to LC3B-decorated vesicles recruited to *S. flexneri*. Images shown correspond to a slice of 2.6 µm thickness. See also Movie S1. Scale bar, 1 μm.

To study the dynamics of septin – LC3B interactions at high resolution, we performed time-lapse Airyscan confocal microscopy of HeLa cells producing GFP-SEPT6 and mCherry-LC3B infected with *S. flexneri* (Fig. S6A). Live cell imaging revealed that septins interact with LC3B-positive membranes during autophagy of septin-caged bacterial cells (Fig. S6A, white arrow, Movie S4). In this case, septins assemble as ∼0.6 µm ring-like structures surrounding LC3B-positive membranes (Fig. S6A, S6B, Movie S5), consistent with septin -Atg8 interactions during autophagosome formation in *Saccharomyces cerevisiae* (Barve et al., 2018). Recent work using different cell types has shown a role for septins during autophagosome formation that depends on direct septin – LC3B interactions (Barve et al., 2018; Toth et al., 2022). Considering this, we hypothesized that septins may also bind to LC3B during autophagy of caged *S. flexneri*. To test this, we performed co-immunoprecipitation assays using GFP-LC3B producing HeLa cells, however, under the conditions tested we could not detect septin – LC3B binding (data not shown). Together, cryo-SXT and live cell imaging data suggest an interaction between septins (well known as membrane-interacting proteins) and autophagic membranes during autophagy of entrapped *S. flexneri* cells.

### K63-linked ubiquitin chains decorate septin cage-entrapped *S. flexneri* for targeting to autophagy

How septin cage-entrapped *S. flexneri* are recognized by autophagy is poorly understood (Mostowy et al., 2010; Mostowy et al., 2011). K63 and K48 polyubiquitin chains are responsible for targeting cargos to autophagic or proteasomal degradation, respectively (Akutsu et al., 2016). During xenophagy, ubiquitin is recognized by autophagy adaptor proteins that also bind LC3B. To further explore how septin-caged *S. flexneri* are targeted to autophagy, we tested if entrapped bacteria are modified by K63 or K48 polyubiquitin chains. We infected HeLa cells producing mCherry-LC3B with *S. flexneri* and quantified the percentage of septin caged bacteria co-localizing with K63 or K48 polyubiquitin (Fig. 4A). Consistent with a role for K63 chains (and not K48 chains) in xenophagy, 49.9 ± 6.9 % of septin cage entrapped *S. flexneri* co-localize with both K63 polyubiquitin and LC3B (Fig. 4B), while only 14.5 ± 4.1% of caged bacteria co-localize with both K48 polyubiquitin and LC3B (Fig. 4C). In support of a role for K63 in promoting the recruitment of LC3B, only 0.8 ± 0.8% of septin caged *S. flexneri* co-localize with LC3B but not K63 polyubiquitin (Fig. 4B). In agreement with a role for septin cages in targeting bacteria to xenophagy, infection of GFP-LC3B-producing HeLa cells showed that 54 ± 10% of GFP-LC3B-positive *S. flexneri* also co-localize with ubiquitin and septins (Fig. 4D).

**Figure 4.**
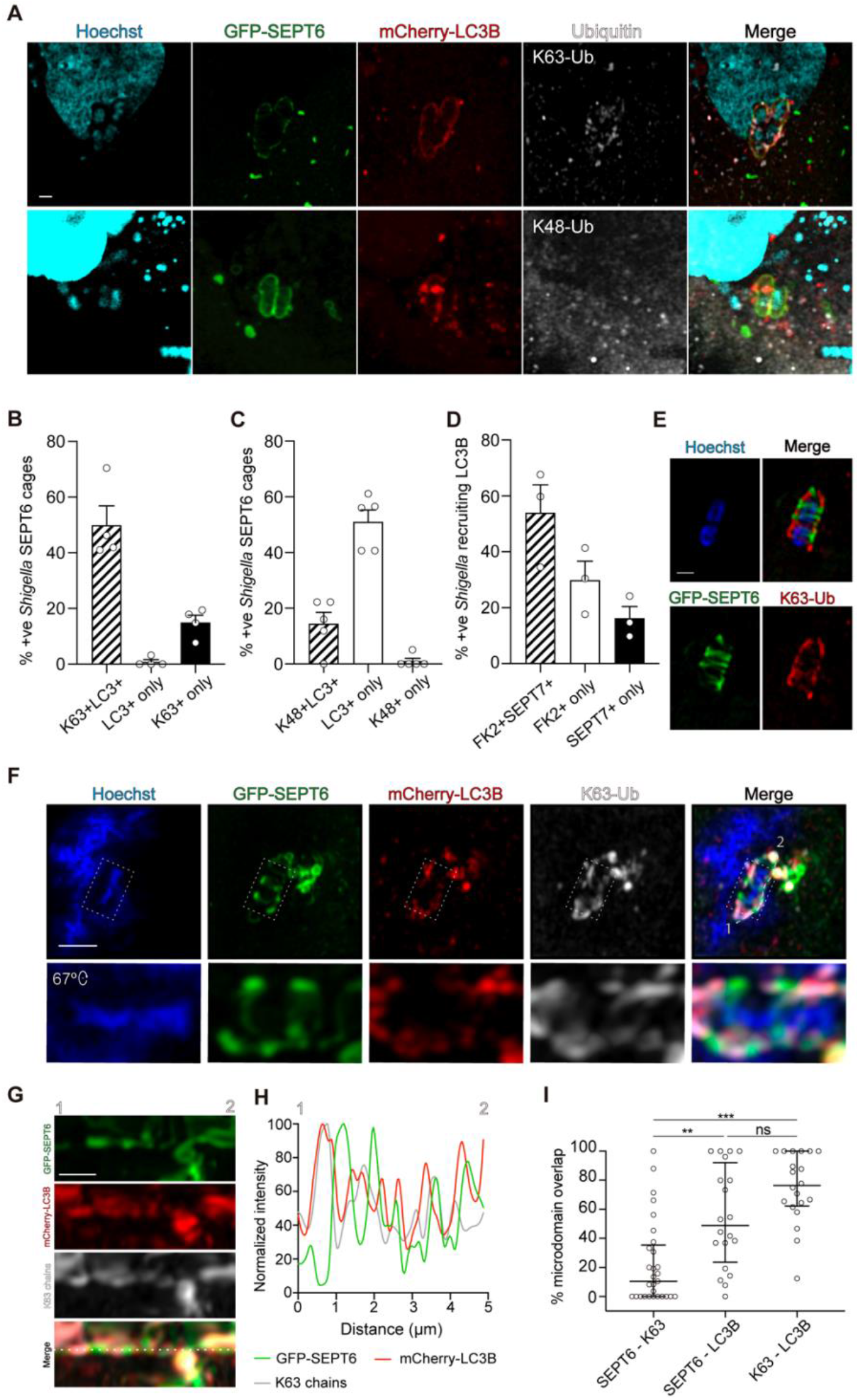
K63-linked ubiquitin chains target *S. flexneri* to autophagy. **(A)** Airyscan confocal image showing septin caged *S. flexneri* co-localizing with LC3B and K63 polyubiquitin (top panel) and septin caged *S. flexneri* co-localizing with LC3B but not K48 polyubiquitin (bottom panel). Scale bar, 1 μm. **(B)** Quantification of *S. flexneri* septin cages co-localizing with LC3B and/or K63 polyubiquitin. Data represents the mean ± SEM from n = 66 *S. flexneri* -septin cages distributed in 4 independent experiments. ***, p < 0.001 by one-way ANOVA and Tukey’
ss post-test. **(C)** Quantification of *S. flexneri* septin cages co-localizing with K48 chains. Data represents the mean ± SEM from n = 191 *S. flexneri* septin cages distributed in 5 independent experiments. ns, p > 0.05 by one-way ANOVA and Tukey′s post-test. **(D)** Quantification of *S. flexneri* decorated with GFP-LC3B co-localizing with ubiquitin (FK2) and/or SEPT7. Data represents the mean ± SEM from n = 149 bacteria distributed in 3 independent experiments. **(E)** Airyscan confocal images showing the formation of separate microdomains of GFP-SEPT6 and K63 polyubiquitin on the surface of *S. flexneri*. Scale bar, 1 μm. **(F)** Airyscan confocal images showing the formation of separate microdomains of septins, K63 polyubiquitin and LC3B on the surface of *S. flexneri*. mCherry-LC3B partially co-localize with both GFP-SEPT6 and K63 chains. Scale bar, 2 μm. **(G)** Representation of the GFP-SEPT6, K63 polyubiquitin and mCherry-LC3B microdomains of the septin cage from panel (E). 1 and 2 mark the beginning and end of the dashed white line from (F). Scale bar, 1 µm. **(H)** Fluorescence intensity profiles of GFP-SEPT6, K63 polyubiquitin and mCherry-LC3B across the white line from panel (G). **(I)** Quantification of the co-localization between GFP-SEPT6, K63 polyubiquitin and mCherry-LC3B. Data represents the median ± interquartile range from n = 29 (GFP-SEPT6 and K63 polyubiquitin), n = 20 (K63 chains and mCherry-LC3B) and n = 20 (GFP-SEPT6 and mCherry-LC3B) bacteria distributed in 3 independent experiments. *, p < 0.001; ***, p < 0.0001; ns, non-significant by Kruskal-Wallis with Dunn′s post-test.

In addition to septin cage entrapment, the host cell can restrict actin-based motility of *S. flexneri* by decorating cytosolic bacteria with guanylate-binding proteins (GBPs) (Li et al., 2017;Piro et al., 2017; Wandel et al., 2017). However, *S. flexneri* can escape from GBP recognition by secreting an E3-ubiquitin ligase (IpaH9.8) that targets GBPs with K48 polyubiquitin for proteasomal degradation (Li et al., 2017; Wandel et al., 2017). Consistent with this, the mutant *S. flexneri* Δ*ipaH9*.*8* cannot ubiquitinate GBPs and is recognized by GBPs more frequently than WT bacteria (Li et al., 2017; Wandel et al., 2017). Considering this, we questioned whether *S. flexneri* can target septins for proteasomal degradation in order to escape from cage entrapment. To test this, we infected HeLa cells with *S. flexneri* WT or the E3 ligase mutants Δ*ipaH1*.*4*, Δ*ipaH2*.*5* or Δ*mxiE* (a strain lacking a transcription factor responsible for the upregulation of many effector genes, including 12 *S. flexneri* -encoded E3 ligases) and quantified the percentage of SEPT7 cages. From these experiments, we did not observe significant differences between *S. flexneri* WT (11.5 ± 0.8%), Δ*ipaH1*.*4* (12.5 ± 0.9%), Δ*ipaH2*.*5* (10.7 ± 1.4%) and Δ*mxiE* (12.8 ± 0.9%), indicating that the E3 ligases tested here do not enable escape from septin cage entrapment (Fig. S7). We do not rule out the possibility of mechanistic redundancy among *S. flexneri* E3 ligases, yet these observations are in agreement with previous work showing that proteasome inhibition does not alter the percentage of septin caged *S. flexneri* (Mostowy et al., 2010).

### K63-linked ubiquitin chains and septins localize to non-overlapping microdomains around *S. flexneri*

To test if septins can promote the ubiquitination of cytosolic *S. flexneri*, we depleted SEPT7 by small interfering RNA (siRNA) (Fig. S8A, S8B) and quantified the percentage of total cytosolic bacteria decorated with total ubiquitin (FK2). Analysis by confocal microscopy revealed no significant difference in the percentage of ubiquitin-positive *S. flexneri* between control (6.2 ± 0.9%) and SEPT7 (5.8 ± 0.4%) siRNA-treated cells (Fig. S8C, S8D). We observed a similar recruitment of K63 polyubiquitin to *S. flexneri* in the case of control (6.2 ± 1.2%) and SEPT7 (5.8 ± 1.2%) siRNA-treated cells (Fig. S8E). These data suggest that recruitment of K63 chains to *S. flexneri* is independent from the recruitment of SEPT7. To test this, we visualized ubiquitinated *S. flexneri* (K63 chains) entrapped in septin cages during the infection of HeLa cells producing GFP-SEPT6 and mCherry-LC3B. Strikingly, Airyscan confocal microscopy showed that GFP-SEPT6 and K63 polyubiquitin form non-overlapping microdomains around the bacterial membrane (Fig. 4E, S9). We then visualized the distribution of GFP-SEPT6, K63 chains and mCherry-LC3B on cytosolic *S. flexneri*. While SEPT6 and K63 polyubiquitin form different microdomains on the bacterial membrane, LC3B partially co-localized with both SEPT6 and K63 polyubiquitin (Fig. 4F, 4G, 4H). We quantified the percentage of GFP-SEPT6, K63 polyubiquitin and mCherry-LC3B co-localizing with each other. In agreement with septins and K63 polyubiquitin recognizing separate *S. flexneri* microdomains, SEPT6 rarely (median 10.5 %) co-localized with K63 chains (Fig. 4I). Consistent with their role in targeting substrates to autophagy, K63 polyubiquitin frequently (median 76.4 %) co-localized with LC3B. Surprisingly, LC3B co-localized with both SEPT6 and K63 polyubiquitin equally (Fig. 4I). Together, these data demonstrate that SEPT6 and K63 polyubiquitin form separate bacterial microdomains during autophagy of septin caged entrapped *S. flexneri*.

## Discussion

The septin cage was first described over 10 years ago (Mostowy et al., 2010; Robertin and Mostowy, 2020). In the case of *S. flexneri*, septins have been shown to bind bacterial membrane for cage entrapment *in vitro* (Lobato-Márquez et al., 2021). Despite intensive research, septin cages had not yet been imaged in their native state inside infected cells, and the interplay between septins and autophagy was unknown. Here, we used correlative light and cryo-SXT to visualize septin cage-entrapped bacterial cells at the nanometer scale to understand their link with autophagosome formation. We discovered that septin cages are strongly correlated with X-ray dense structures surrounding *S. flexneri*. In agreement with a role for mitochondria during cage entrapment of *S. flexneri*, cryo-SXT showed that mitochondria are tightly bound to septin cages *in situ*. We also showed that septin caged *S. flexneri* are decorated with K63 polyubiquitin (via a process separate from recruitment of septins) to target entrapped bacteria to autophagy. Airyscan confocal microscopy and correlative light and cryo-SXT suggest that septins interact with LC3B-decorated membranes during autophagy of *S. flexneri*. Taken together, a model emerges, where: i) septins recognize poles of cytosolic *S. flexneri* for cage entrapment; ii) an unknown E3-ubiquitin ligase/s decorates separate regions of the bacterial membrane (not covered by septins) with K63 polyubiquitin; iii) autophagy adaptor proteins link ubiquitinated bacteria to LC3B; iv) septins interact with LC3B-positive membranes during autophagy of *S. flexneri*. This process ultimately leads to the encapsulation of septin cage entrapped *S. flexneri* into autophagosomes targeted to lysosomal fusion.

Our data show that the host cell ubiquitin machinery targets microdomains on the bacterial membrane distinct from the microdomains bound to septins. Ubiquitination is a sequential enzymatic cascade mediated by ubiquitin-activating (E1), ubiquitin-conjugating (E2), and ubiquitin-ligating (E3) enzymes (Hershko and Ciechanover, 1998). In the human genome, there are 2 E1 genes, 30 – 50 E2 genes and > 600 E3-encoding genes (Zheng and Shabek, 2017); this diversity has made it highly challenging to identify the specific enzyme/s targeting *S. flexneri*. The E3-ubiquitin ligase LRSAM1 has been shown to decorate a *S. flexneri* Δ*icsB* mutant (IcsB is an effector correlated with autophagy and septin cage avoidance) with K63 and K27 polyubiquitin (Huett et al., 2012; Mostowy et al., 2010). For this study, we tried to ubiquitinate *S. flexneri in vitro* using purified E1, UBE2D2 and LRSAM1, but under the conditions tested we could not detect ubiquitinated *S. flexneri* (data not shown). As far as we know, the E3-ubiquitin ligase/s that decorate *S. flexneri* WT with K63 polyubiquitin have not yet been identified. The E3 ligase RNF213 has recently been shown to ubiquitinate Lipid A on the outer membrane of *S*. Typhimurium (Otten et al., 2021). Considering the structure of Lipid A is conserved between *S. flexneri* and *S*. Typhimurium, it is tempting to speculate that RNF213 may also ubiquitinate *S. flexneri*. As both septins (Lobato-Márquez et al., 2021) and RNF213 (Otten et al., 2021) target the bacterial membrane, this could explain the distinct microdomains we observed during *S. flexneri* infection. New technologies in cellular microbiology (López-Jiménez and Mostowy, 2021), such as proximity-dependent biotinylation coupled with mass spectrometry (Liu et al., 2020), may prove useful to identify the specific E3-ubiquitin ligase/s targeting *S. flexneri*. Alternatively, high-throughput microscopy combined with CRISPR-Cas libraries has been used to identify the E3-ubiquitin ligases involved in *S*. Typhimurium ubiquitination (Heath et al., 2016; Polajnar et al., 2017). Post-translational modifications are well known to play an important role in host-pathogen interactions (Chambers and Scheck, 2020). Septins can be post-translationally modified by ubiquitination, SUMOylation, acetylation and phosphorylation (Hernandez-Rodriguez and Momany, 2012). We recently demonstrated that purified septin proteins can recognize bacterial membranes in the absence of additional host cell factors, including post-translational modifications (Lobato-Márquez et al., 2021). Together with data showing that K63 polyubiquitin does not co-localize with septins at the *S. flexneri* septin cage, we propose that K63 polyubiquitin is not required for septin cage assembly. Considering that SUMOylation has been shown to regulate septin assembly during cytokinesis (Hernandez-Rodriguez and Momany, 2012; Ribet et al., 2017), it is next of great interest to study the role of post-translational modifications (including SUMO and ubiquitin linkages other than K63 polyubiquitin) during septin-mediated cell-autonomous immunity.

Cryo-SXT is a powerful imaging technique used to study membrane-based cellular organelles, including autophagy. Here, we use correlative light and cryo-SXT to study septin-autophagy interactions during *S. flexneri* cage entrapment. During autophagy of *S. flexneri*, we show that septins interact with LC3B-positive membrane. Unfortunately, the resolution obtained in our study using cryo-SXT did not permit the visualization of individual septin filaments, nor how they interact with autophagic membranes, and thus the precise role of septin– LC3B interactions during autophagy of *S. flexneri* requires further investigation. Considering the recent visualization of septin filaments on septin cages reconstituted *in vitro* (Lobato-Márquez et al., 2021), and the combination of focused ion beam milling with cryo-electron tomography (Wagner et al., 2020), it will be important to explore *in situ* how septins organize on autophagic membrane at nanometer resolution. In parallel, combining purified septin proteins with autophagosomes reconstituted *in vitro* may illuminate the molecular mechanisms underlying septin – autophagosome interactions (Chang et al., 2021; Sawa-Makarska et al., 2020). In this way, an in depth understanding of septin – autophagy interactions can help to identify novel approaches for bacterial infection control.

## Supporting information

ImageJ macros used

## Author contribution

D.L.-M., J.J.C. and S.M. conceived the study. D.L.-M., J.J.C., A.T.L.-J., M.E.D., J.N.P. and S.M. analyzed and interpreted the data. D.L.-M. and J.J.C. performed and analyzed cryo-SXT experiments. D.L.-M. and A.T.L-J. performed all fluorescence microscopy experiments. D.L.-M., M.E.D. and J.N.P. performed and analyzed *in vitro* ubiquitination assays. D.L.-M. and S.M. wrote the manuscript; all authors commented on the manuscript.

## Acknowledgments

We thank Mostowy lab members for helpful discussions. We thank Paul Simpson and the Imperial College EM facility, and the Centro Nacional de Biotecnología EM facility for their help with the vitrification of samples. We thank David C. Rubinsztein for the GFP-LC3-producing HeLa cell line. We thank Sharon Tooze for the plasmid encoding *mCherry-LC3B*. We thank Felix Randow for the *S. flexneri* Δ*mxiE* strain. D.L.-M. was funded by the European Union’s Horizon 2020 research and innovation program under the Marie Skłodowska-Curie grant agreement No. H2020-MSCA-IF-2016-752022 (INCAGE), and the ALBA synchrotron (grant No. 2018093019). A.T.L-J. is funded by a Swiss National Science Foundation Early Postdoc.Mobility Fellowship (grant No. P2GEP3_188277) and the European Union’s Horizon 2020 research and innovation program under the Marie Skłodowska-Curie grant agreement No. H2020-MSCA-IF-2020-895330. Work in the J.N.P laboratory is supported by the National Institute of General Medical Sciences (R35GM142486). Work in the S.M. laboratory is supported by a European Research Council Consolidator Grant (grant agreement No. 772853-ENTRAPMENT), Wellcome Trust Senior Research Fellowship (206444/Z/17/Z) and the Lister Institute of Preventive Medicine.

## Material and Methods

### Reagents

The following antibodies were used: rabbit anti-SEPT7 (#18991, IBL), rabbit anti-ubiquitin Lys-63-specific (#05-1308, Merck), rabbit anti-ubiquitin Lys-48-specific (#05-1307, Merck), mouse FK2 (#PW8810, Enzo Life Sciences), Alexa-555-conjugated anti-rabbit antibody (#10082602, ThermoFisher Scientific), Alexa-647-conjugated anti-rabbit antibody (#A27040, ThermoFisher Scientific). Hoechst (#H3570, ThermoFisher Scientific) was used through the manuscript to stain for *S. flexneri* and epithelial cell DNA.

### Bacterial strains and culture conditions

Unless otherwise indicated, *Shigella flexneri* 5a str. M90T producing the adhesin AfaI (Mostowy et al., 2010) was used throughout the manuscript. *S. flexneri* was grown in trypticase soy broth (TCS)-agar containing 0.01% (w/v) congo red to select for red colonies, indicative of a functional T3SS. Conical polypropylene tubes (#CLS430828, Corning) containing 5 ml of TCS were inoculated with individual red colonies of *S. flexneri* and were grown ∼16 h at 37 ºC with shaking at 200 rpm. The following day, bacterial cultures were diluted in fresh pre-warmed TCS (1: 50 v/v), and cultured until an optical density (OD, measured at 600nm) of 0.6. To maintain the plasmid encoding *afaE* (AfaI-encoding gene), TCS was supplemented with 100 µg/ml of carbenicillin.

*Escherichia coli* strains were grown in Lysogeny-Broth (LB) in conical polypropylene tubes at 37 °C with shaking at 220 rpm. *E. coli* DH5α was used to purify pKD46 and pKD4 plasmids (Datsenko and Wanner, 2000), and LB was supplemented with 100 μg/ml of carbenicillin or 50 μg/ml of kanamycin, respectively.

Bacterial stocks were stored in 10% glycerol at -80 ºC.

### Design of bacterial mutant strains

Primers used in this study were designed using Benchling (https://benchling.com) and are listed in Supplementary Table 1. *S. flexneri* mutants were engineered using λ-Red-mediated recombination (Datsenko and Wanner, 2000). In brief, kanamycin resistance-encoding DNA cassettes were amplified using pKD4 plasmid as template and primers containing 50 bp nucleotides homologous to the site of insertion. Resulting DNA fragments were electroporated in *S. flexneri* electrocompetent cells producing λ-Red recombinase and plated in TSA plates supplemented with 0.01% of congo red and 50 μg/ml of kanamycin. All strains were verified by PCR.

### Mammalian cell culture

HeLa (ATCC CCL-2) cells were grown at 37 ºC and 5% CO_2_ in Dulbecco’s Modified Eagle Medium (DMEM, GIBCO) supplemented with 10% fetal bovine serum (FBS, Sigma-Aldrich). GFP–SEPT6 producing HeLa cells (Sirianni et al., 2016) were grown as above in DMEM supplemented with 10% FBS and 2 µg/ml of puromycin. GFP–LC3B producing HeLa cells (Runwal et al., 2019) were grown as above in DMEM supplemented with 10% FBS.

### HeLa cell plasmid transfection

8 × 10^4^ HeLa cells were seeded in 6-well plates (Thermo Scientific) containing 22 × 22 mm glass coverslips, Quantifoil (R2/2, Quantifoil Micro Tools) Au-EM finder grids, coated with holey carbon, or MatTek dishes 2 days before transfection. Plasmid transfections were performed in 1 ml DMEM with 250 ng of a plasmid encoding *mCherry-LC3B* (gift from Sharon Tooze lab) using JetPEI (Polyplus transfection) as described in (Mazon Moya et al., 2014) and incubated at 37 ºC and 5% CO_2_ for 6 h. Then, the medium was substituted with 2 ml of fresh pre-warmed DMEM supplemented with 10% FBS until the following day when cells were infected.

### Infection of human cells

For experiments involving paraformaldehyde (PFA)-fixed samples (Fig. 1C, Fig. 4, Fig. S3, Fig. S7 and Fig. S8) 9 × 10^4^ HeLa cells were seeded in 6-well plates (Thermo Scientific) containing 22 × 22 mm glass coverslips 2 days before the infection. For experiments involving live cell imaging (Fig. S5 and Fig. S6) 9 × 10^4^ HeLa cells were seeded in MatTek glass-bottom dishes (MatTek corporation). For experiments involving correlative light and cryo-SXT (Fig. 1, Fig. 2, Fig. 3, Fig. S1 and Fig. S4) 8 × 10^4^ HeLa cells were seeded in 6-well plates containing or Quantifoil Au-EM finder grids. Bacterial cultures were grown as described above, and cell cultures were infected with *S. flexneri afaE* at a multiplicity of infection (MOI, bacteria: cell) of 10: 1. Then, plates were placed at 37 ºC and 5% CO_2_ for 30 min. Infected cultures were washed 2X with phosphate buffered saline (PBS) pH 7.4 and incubated with fresh DMEM containing 10% FBS and 50 mg/mL gentamicin at 37 ºC and 5% CO_2_ up to 3 h. For live cell imaging experiments, DMEM was substituted by OPTI-MEM containing 10% FBS and 50 mg/mL gentamicin and MatTek dishes placed on an Airyscan 880 confocal microscope coupled to a temperature-controlled incubator (37 ºC).

### Cryo-epifluorescence microscopy

After infection, HeLa cells were fixed by plunge-freezing using a Vitrobot Mark IV (Thermo Fisher). Prior to vitrification, samples were slightly fixed with 1% PFA for 5 min to inactivate bacteria and imaged using an Axiovert Z1 driven by ZEN Blue 2.3 software (Carl Zeiss) epi-fluorescence microscope or a confocal microscope LSM710 (Carl Zeiss) driven by ZEN 2010 software. To stain for mitochondria, samples were incubated with 100 nM of Mitotracker Red (Invitrogen) for 30 min before vitrification. In all cases, grids were incubated with 100 nm fiducial gold nanoparticles (that would help during the alignment of the tilt series) for 30 s before vitrification. Following vitrification, grids were shipped to the Mistral beamline at the ALBA synchrotron (Barcelona, Spain). Vitrified grids were then transferred in liquid nitrogen to the cryo-correlative cooling stage Linkam CSM196 (Linkam Scientific Instruments) to hold samples at a stable -190°C during analysis. The cryo-stage was inserted into an AxioScope A1 (Carl Zeiss) epifluorescence microscope with a N-Achroplan 50x/0.6 Ph1 objective and imaged with a CCD AxioCam ICm1 (Carl Zeiss).

Cryo-fluorescence microscopy was used to pre-select vitrified samples and map the position of cells. Selected samples were then transferred to the Mistral synchrotron beamline at liquid nitrogen temperature.

### Soft X-ray cryo-tomography

Grids were visualized on-line with a visible light microscope integrated within the X-ray microscope to correlate cell positions identified with epifluorescence and cryo-epifluorescence images prior to sample loading into the Mistral beamline. Zero-degree soft X-ray projection mosaics were acquired to image the areas of interest and define the tomogram acquisition areas. Tilted-series were acquired at 520 eV photon energy from -70º to 70º for each degree, using a 25-nm zone plate. The exposure time used was associated to sample conditions (thickness), and ranged from 1 to 4 sec. Pixel size was set to 11.5 nm.

Tilted series were normalized to the flatfield using the XMIPP 3 software package (de la Rosa-Trevin et al., 2013), aligned with IMOD (Kremer et al., 1996) and reconstructed with the TOMO3D software SIRT algorithm using 30 iterations (Simultaneous Iterative Reconstructive Technique) (Agulleiro and Fernandez, 2011). Semiautomatic segmentation of volumes was carried out with Amira software (ThermoFisher Scientific), and volumes were represented with Chimera software (Pettersen et al., 2004).

### Epifluorescence microscopy of infected cells

HeLa cells producing stably GFP-SEPT6 seeded in MaTtek dishes and transfected with a plasmid encoding mCherry-LC3B (as described above) were infected with *S. flexneri* AfaI for 30 min at 37ºC and 5% CO_2_. Samples were then transferred to a temperature-controlled chamber (37ºC and 5% CO_2_) and imaged in FluoroBrite medium (Life Technologies) supplied with 5% FCS, 4 mM L-glutamine and 50 µg/ml gentamicin. Epifluorescence imaging was performed using an Axiovert Z1 microscope driven by ZEN Blue 2.3 software. Microscopy images were obtained using z-stack image series taking 11 slices.

### *In vitro* reconstitution of *S. flexneri* -septin cages

*In vitro* reconstitution of septin cages was performed as described in Lobato-Márquez et al., 2021. *S. flexneri* culture was grown 16 h in conical polypropylene tubes containing 5 ml of M9-Tris (50 mM Tris-HCl pH8, 50 mM KCl, 0.5 mM MgCl_2_, 0.1 mM CaCl_2_, 1 mM MgSO_4_) salts supplemented with a mix of nutrients (45 µg/ml L-methionine, 20 µg/ml L-tryptophan, 12.5 µg/ml nicotinic acid, 10 µg/ml vitamin B1, 1% glucose, 0.5 % casein hydrolysate, 0.1 % fatty acid-free BSA) -M9-Tris-CAA-at 37 ºC with shaking at 200 rpm. The following day, bacterial cultures were diluted in 10 ml of fresh pre-warmed M9-Tris-CAA (1: 100 v/v) in conical polypropylene tubes and cultured until an OD_600_ of 0.6. 1.2 ml of bacterial cultures were centrifuged in Low Protein Binding tubes (ThermoFisher Scientific) at 800 × g for 2 min at RT and the supernatant was removed. Purified recombinant septin (SEPT2/msGFP– SEPT6/SEPT7) hetero-oligomers in septin storage buffer (50 mM Tris pH8, 300 mM KCl, 5 mM MgCl_2_ and 5 mM DTT) were thawed on ice, diluted, and added to the *in vitro* reconstitution solution at a final concentration of 240 nM (yielding a final buffer composition of 50 mM Tris pH8, 50 mM KCl, 0.5 mM MgCl_2_ and 1 mM DTT). Low Protein Binding tubes containing the bacteria in the *in vitro* reconstitution solution were placed in opaque conical polypropylene tubes and incubated at 37 ºC with shaking at 220 rpm for 2 h. Following the *in vitro* reconstitution reaction, samples were immediately placed on ice. To remove unbound recombinant septin hetero-oligomers, samples were centrifuged at 800 × g at 4 ºC for 1.5 min. Supernatant was carefully removed, bacterial pellet containing bound septins resuspended in 300 µl of ice-chilled M9-Tris-CAA buffer and centrifuged at 800 × g at 4 ºC for 2 min. This step was repeated one more time, to ensure removal of unbound septins, and pellets were finally resuspended in 100 µl of ice-chilled M9-Tris-CAA buffer. We placed 3 µl of this *in vitro* reconstitution reaction on EM grids that were plunge frozen and imaged by correlative light and cryo-SXT as described above.

### Immunostaining and confocal microscopy

Infected cells were washed 3X with PBS pH 7.4 and fixed 15 min in 4% PFA (in PBS) at room temperature. Fixed cells were washed 3X with PBS pH 7.4 and subsequently permeabilized 5 min with 0.1% Triton X-100 (in PBS). Cells were then washed 3 − 6X in PBS and incubated 1 h 30 min with primary anti-SEPT7, anti-FK2, anti-K63-Ub or anti-K48-Ub antibody diluted in PBS supplemented with 0.1% Triton X-100 and 1% bovine serum albumin. Note that HeLa cells produce SEPT6 and SEPT7, that together with SEPT2 and SEPT9, produce a hetero-oligomer SEPT2-SEPT6-SEPT7-SEPT9 that assembles into filaments and higher-order structures (Kim et al., 2011; Mostowy and Cossart, 2012). SEPT7 is a septin essential for heter-oligomer formation, and therefore SEPT7 staining represent the cellular distribution of the other septins (Martins et al., 2022). Cells were then washed 3 − 6X in PBS and incubated 45 min with Alexa-555-conjugated anti-rabbit or Alexa-647-conjugated anti-rabbit secondary antibody diluted 0.1% Triton X-100 (in PBS). Cells were then washed 3 − 6X in PBS and incubated with a solution of 0.1% Triton X-100 (in PBS) containing Hoechst. Coverslips were placed on glass slides and samples were preserved with aqua polymount mounting medium (ID#18606, Polyscience).

Fluorescence microscopy was performed using a 63x/1.4 C-Plan Apo oil immersion lens on a Zeiss LSM 880 confocal microscope driven by ZEN Black software. Microscopy images were obtained using z-stack image series taking 8 – 16 slices.

For Airyscan confocal live cell imaging MatTek dishes containing infected cells producing GFP-SEPT6 and mCherry-LC3B were placed on a temperature-regulated chamber (37ºC and 5% CO_2_) and imaged using a 63x/1.4 C-Plan Apo oil immersion lens on a Zeiss LSM 880 confocal microscope driven by ZEN Black software. Microscopy images were obtained over time using z-stack image series taking 8 – 16 slices using the Airyscan fast super-resolution (SR) mode.

### Image processing, quantification and statistical analysis

Confocal images of fixed samples were processed using Airyscan processing (Weiner filter) using “Auto Filter” and “3D Processing” options in ZEN Blue software. Confocal images of live samples were processed using Airyscan processing (Weiner filter) using “Auto Filter” and “3D Processing” options in ZEN Black software. Epifluorescence images were deconvolved using ZEN Blue.

Image quantifications were performed in Fiji. Where possible, fluorescence microscopy images were randomized using the macro for Fiji Filename_Randomizer.

Correlation between fluorescence microscopy and 2D soft X-ray images was done using the plugin ec-CLEM for Icy (Institut Pasteur, Paris, France).

Extraction of quantitative information related to mitochondrial anisotropy (deviation from spherical shape) on cryo-SXT volumes was performed using Amira software (ThermoFisher Scientific). Extraction of quantitative information related to mitochondrial branch/rod length on epifluorescence images was performed using the Fiji plugin MiNa (Valente et al., 2017).

The analysis of K63 chains and LC3B associated to *S. flexneri* microdomains was performed in ImageJ. Airyscan confocal slices around the bacterial sagittal plane were maximum projected. Binary masks were then generated for each individual channel and combined. The regions of interests (ROI) were fitted to an ellipse, and its long axis (identifying the bacterial poles) was determined mathematically. ROI were reduced to 0.1 -0.2 μm to fit the peak of maximum intensity of the microdomains and flattened between the points of intersection with the long axis. Finally, the intensity profile of the generated flattened image was quantified. Co-localization between microdomains was calculated defining that a microdomain has a normalized intensity ≥ 60% in the analyzed region.

Statistical analysis was performed in GraphPad Prism (v8.4, La Jolla, USA). Data represent the mean ± standard error of the mean (SEM) or median and interquartile range. Fold changes were calculated from each independent experiment and the mean ± SEM are given in the text. A Student’s t-test (two-tailed), Mann-Whitney test or one-way ANOVA were used to test for statistical significance, with p < 0.05 considered as significant. All statistical details including statistical tests, significance, value of the number of experimental replicates and bacterial cells quantified can be found in the figure legends.

All figures were designed using Adobe Illustrator CC 2018.

## Figures

**Supplementary Figure S1.**
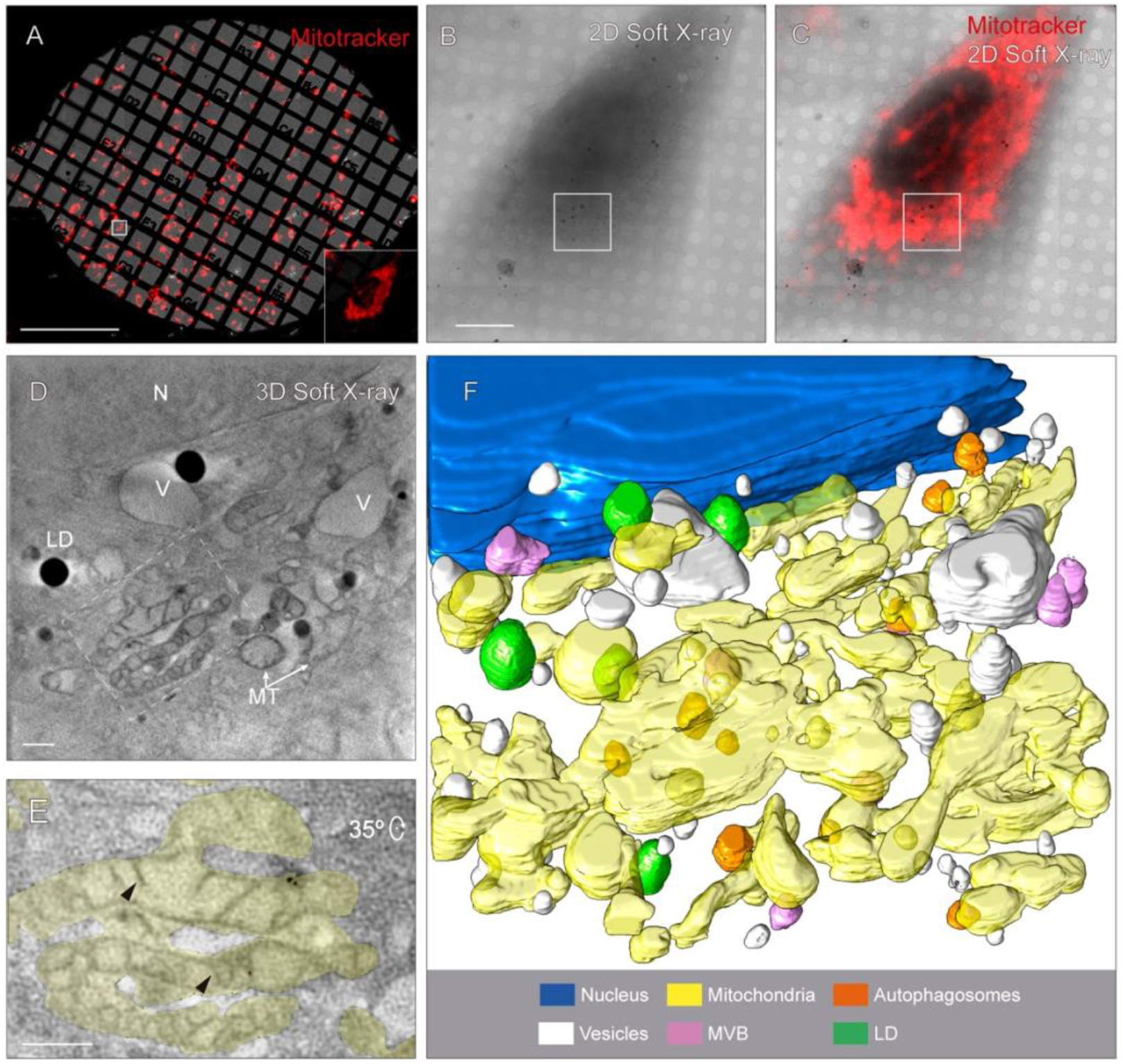
Workflow for correlative light and cryo-SXT. To study the interplay between *S. flenxeri* septin cage entrapment and autophagy, we first developed a correlative light and cryo-SXT pipeline. We seeded human epithelial HeLa cells on Quantifoil (R2/2)-coated Au finder grids and plunge-freezed (vitrified) them in liquified ethane to preserve cellular structures. Before vitrification, cells were stained with Mitotracker Red, and samples were incubated with 100 nm gold fiducials to help with 3D reconstruction of tomograms. Vitrified samples were transferred to a Linkam stage and screened under liquid nitrogen temperature using a cryo-epifluorescence microscope to verify ice thickness, grid quality and locate areas of interest. Selected grids were transferred to a cryo-SXT microscope (operating under liquid nitrogen temperature) where grids were imaged by an on-line cryo-epifluorescence microscope (Fig. S1A). Scale bar, 500 μm. **(B)** Areas of interest (white square on (A) were subsequently imaged by soft X-rays, creating a mosaic that permitted identification of events of interest. Scale bar, 10 μm. **(C)** Correlating fluorescence and X-ray data permitted the identification of regions that could be imaged by X-ray tomography (e.g. white square on (B, C)). **(D)** After cryo-SXT was performed, raw tilt series were aligned and reconstructed. Images shown corresponds to a slice of 3.7 µm thickness. LD, lipid droplet; MT, mitochondria; N, nucleus; V, vesicles. Dotted square represents segmented mitochondria in (E). Scale bar, 1 μm. **(E, F)** Reconstructed tomograms were segmented semi-automatically and rendered in 3D. In the inset of panel (E) it is shown a selected area from (D) depicting ultrastructural details of mitochondria. Mitochondrial crystae are indicated with black arrowheads. This approach enabled visualization of cell volumes at ∼30 nm resolution by soft X-ray and its correlation with fluorescence data. From this pipeline, we could unambiguously identify cell nuclei, mitochondria and lipid droplets, as well as different types of vesicles and autophagosomes (see Fig. S2).

**Supplementary Figure S2.**
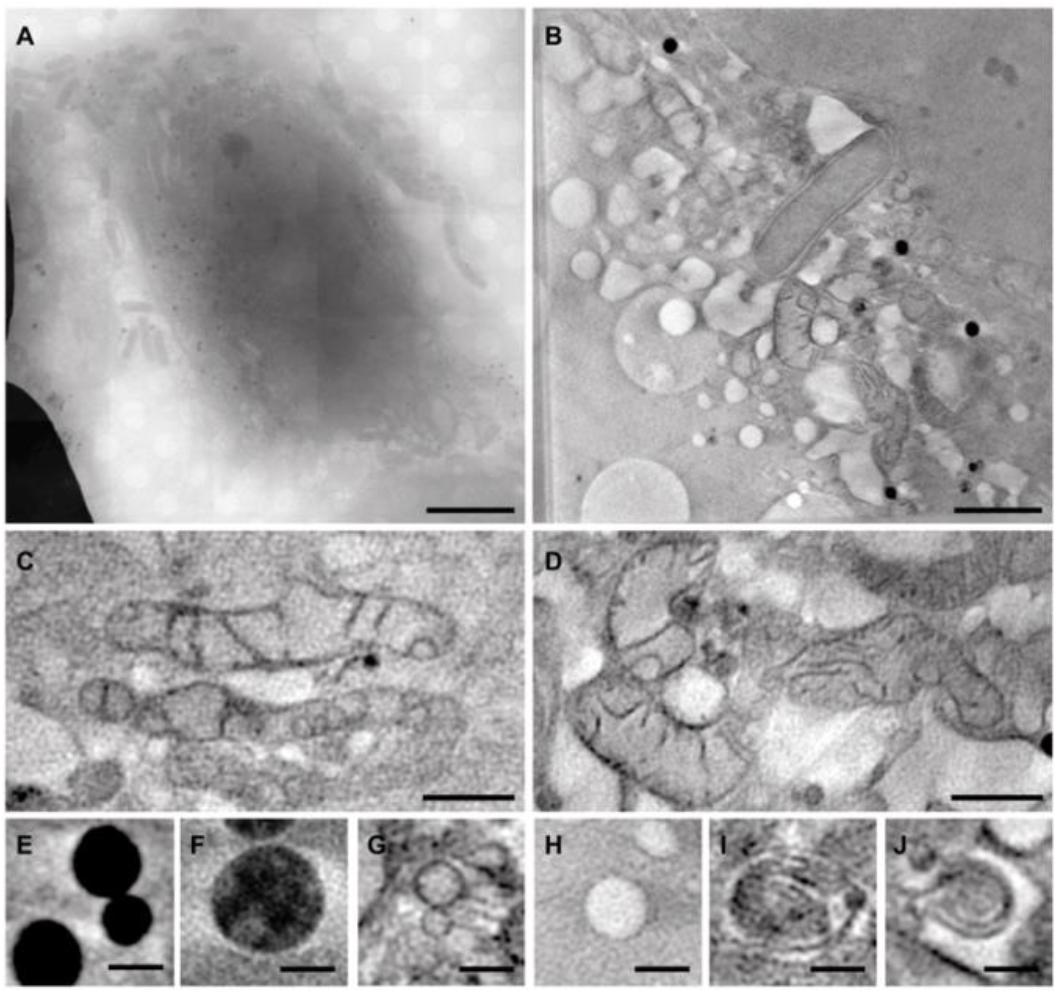
Cryo-SXT atlas of HeLa cells in the context of *S. flexneri* infection. **(A)** Mosaic overview of a *S. flexneri*-infected HeLa cell. Scale bar, 10 μm. **(B-J)** Represent cell volumes containing different type of membrane-based host organelles or bacteria **(B)** Individual *S. flexneri* cell Scale bar, 2 μm. **(C, D)** Mitochondria appear as membranous cellular compartments with a lower X-ray absorbing inner part that contain membranous layers (cristae). Scale bar, 1 μm. (E) Lipid droplets were defined as spherical organelles with very high X-ray absorption due to the enrichment in lipids. Scale bar, 500 nm. **(F)** Multivesicular body, endocytic compartment that contain multiple vesicles inside, are seen as vesicles with variable X-ray absorption densities inside. Scale bar, 500 nm. **(G, H)** Vesicles are observed as endocytic compartments that contain a homogeneous inner X-ray absorption density. Scale bar, 500 nm. **(I, J)** Autophagolysosomes can be identified as endocytic compartments that show concentric membranous structures. Scale bar, 500 nm.

**Supplementary Figure S3 related to Figure 1.**
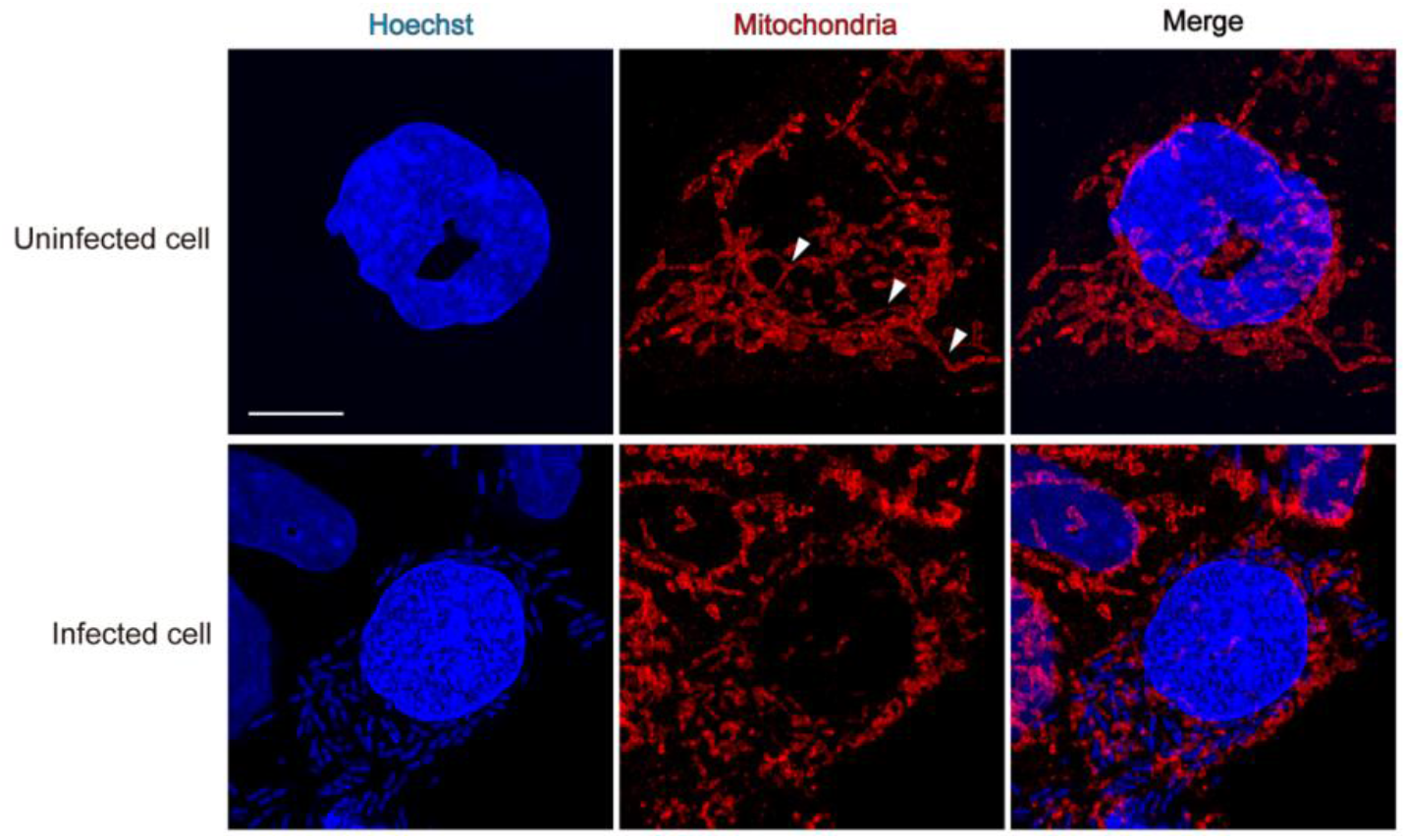
Visualization of mitochondrial dynamics during *Shigella flexneri* infection by Airyscan confocal microscopy. Airyscan confocal images showing elongated mitochondria in the absence of infection (top panels white arrowheads) and mitochondrial fragmentation upon *S. flexneri* infection (bottom panels). Scale bar, 10 μm.

**Supplementary Figure S4 related to Figure 2.**
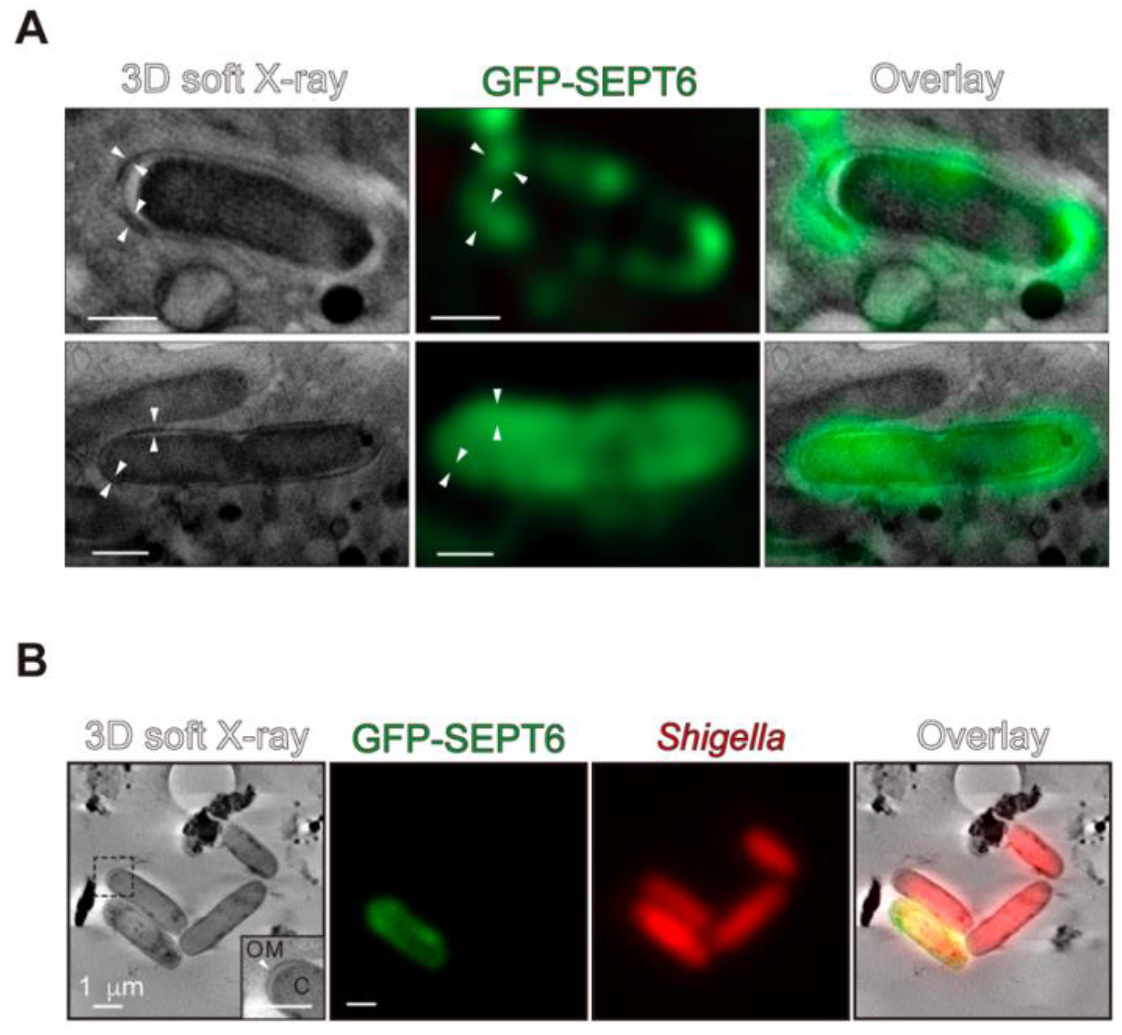
Fluorescent signal of GFP-SEPT6 correlates to increased soft X-ray densities (arrowheads). **(A)**Additional examples related to Figure 2. Scale bar, 1 μm. **(B)** Representative example of a *S. flexneri* cell showing an extended periplasm at the bacterial cell pole, and next to it a separate bacterium entrapped in a septin cage reconstituted *in vitro* using purified septin complexes. Note that in this case there is no additional source of host membrane and proteins and thus septin cages cannot be visualized as X-ray dense structures. Scale bars, 1 μm.

**Supplementary Figure S5 related to Figure 3.**
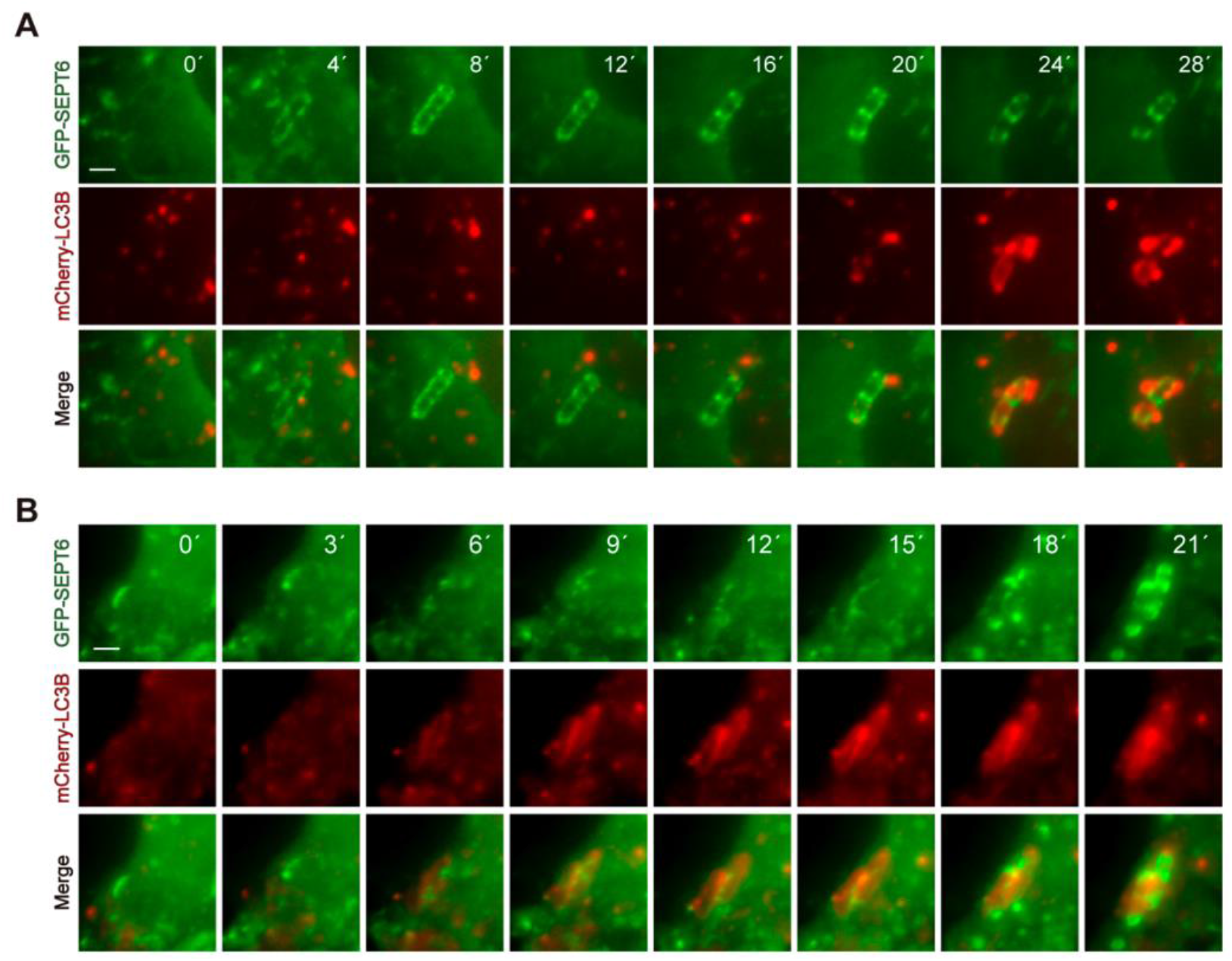
Dynamics of *Shigella* septin cage entrapment and autophagy. **(A)** Epifluorescence time-lapse showing the entrapment of *S. flexneri* in a septin cage (labeled with GFP-SEPT6) followed by autophagosome formation (labeled with mCherry-LC3B). Scale bar, 2 µm. See also Movie S3. **(B)** Epifluorescence time-lapse showing the decoration of *S. flexneri* cells with mCherry-LC3B that are subsequently entrapped in 2 septin cages (labeled with GFP-SEPT6). Scale bar, 2 µm. See also Movie S4.

**Supplementary Figure S6 related to Figure 3.**
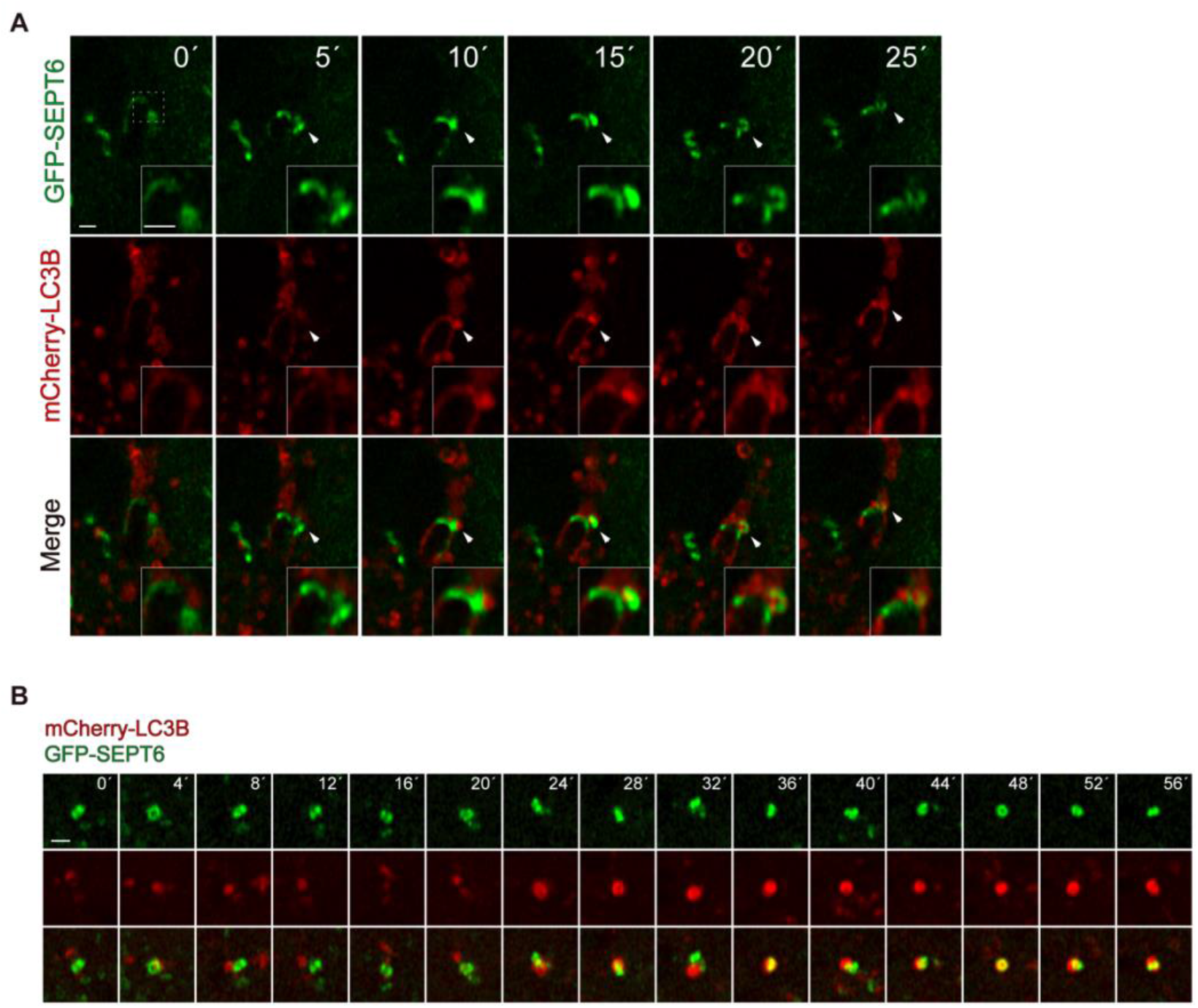
Septins co-localize to autophagosomes as ring-like structures during infection of *S. flexneri*. **(A)** Airyscan fast super-resolution time-lapse showing a caged bacterium, where septins promote the fusion of LC3B to *S. flexneri*. White arrowheads point to the area where septins promote the recruitment of LC3B. See also Movie S5. Scale bar, 1 μm. **(B)** Airyscan fast super-resolution time-lapse showing a septin ring (labeled with GFP-SEPT6) bound to a forming autophagosome (labeled with mCherry-LC3B). See also Movie S6. Scale bar, 1 μm.

**Supplementary Figure S7.**
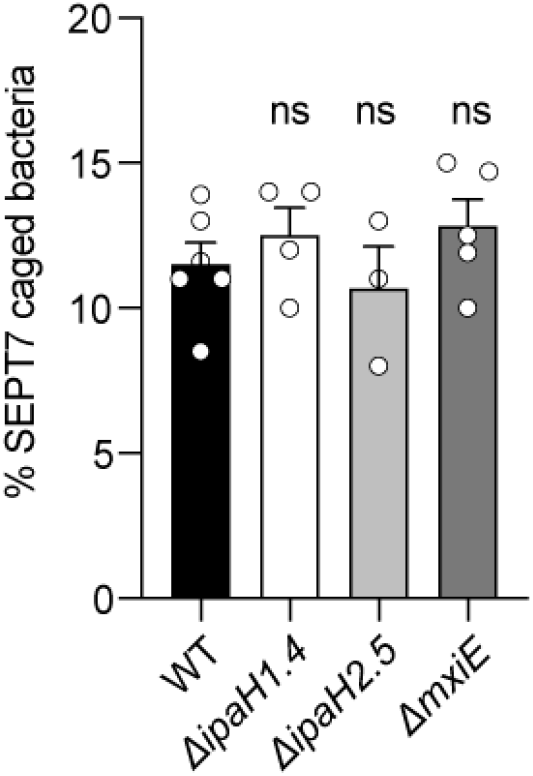
*S. flexneri* E3-ubiquitin ligases do not impact septin cage entrapment. Percentage of *S. flexneri* M90T entrapped in SEPT7 cages in HeLa cells infected for 3h 40 min. Data represents the mean ± SEM from n = 1,466 (WT), n = 821 (Δ*ipaH1*.*4*), n = 851 (Δ*ipaH2*.*5*) and n = 1,089 (Δ*mxiE*) bacterial cells distributed in, at least, 3 independent experiments. ns, p > 0.05 by one-way ANOVA and Tukey’
ss post-test.

**Supplementary Figure S8 related to Figure 4.**
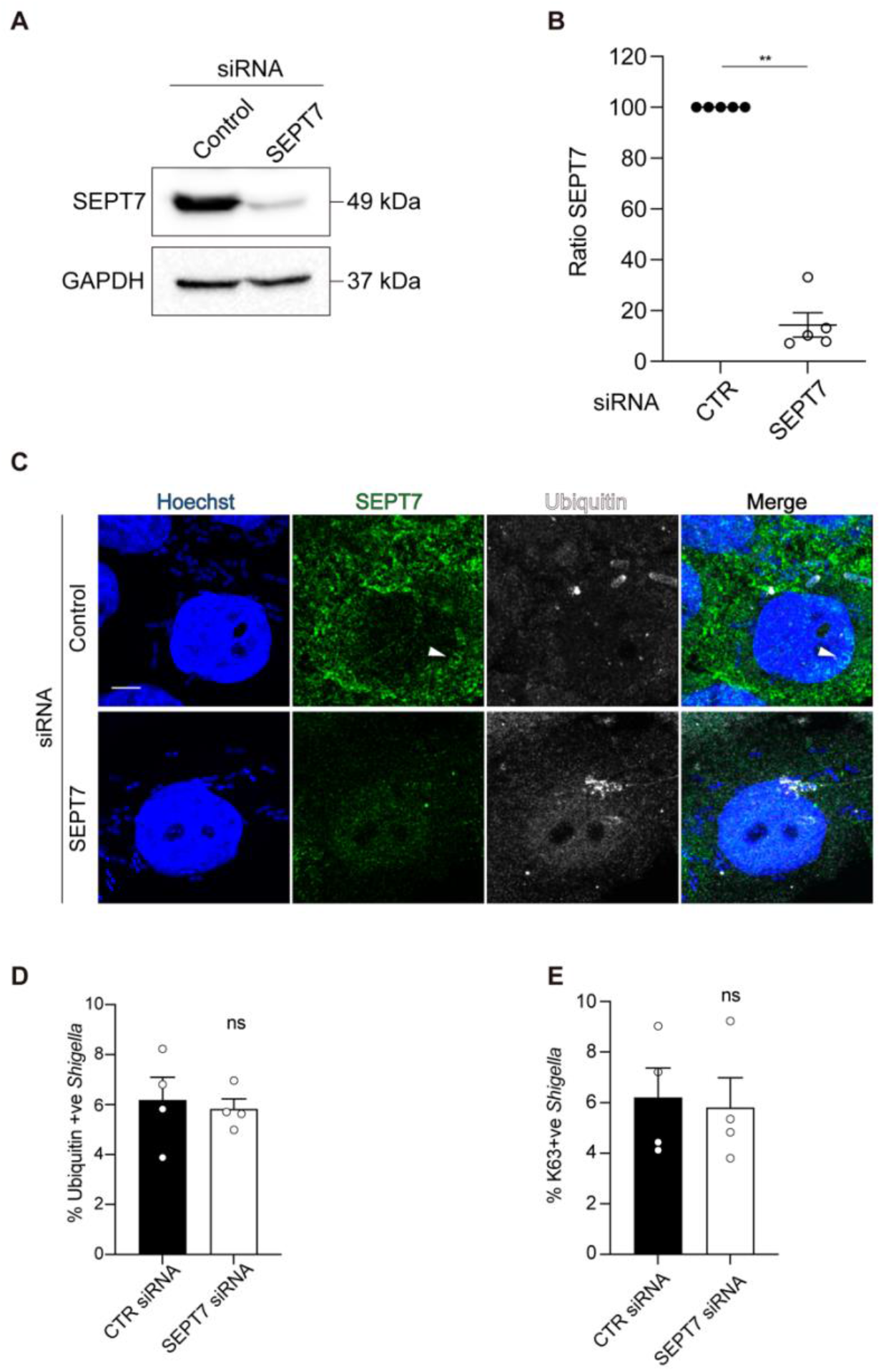
Septins and K63 chains are independently recruited to *S. flexneri*. **(A)** Hela cells were treated with control or SEPT7 siRNA for 72h. Whole cell lysates were immunoblotted for SEPT7 to show the efficiency of depletion. GAPDH was used as loading control. **(B)** Western blot densitometry of 5 independent samples from (A) showing the absence of SEPT7 protein after siRNA treatment. **(C)** Airyscan confocal images showing *S. flexneri* positive with total (FK2) ubiquitin in HeLa cells infected for 3 h and treated with control (top) or SEPT7 (bottom) siRNA. White arrowhead, septin cage. Scale bar, 5 μm. **(D)** Quantification of *S. flexneri* decorated with total ubiquitin in the presence (CTR siRNA) or absence (SEPT7 siRNA) of SEPT7. Data represents the mean ± SEM from n = 1,578 (control siRNA) and n = 1,445 (SEPT7 siRNA) bacteria distributed in 4 independent experiments. ns, p > 0.05 by two-tailed Student’
ss t-test. **(E)** Quantification of *S. flexneri* decorated with K63 polyubiquitin in the presence (CTR siRNA) or absence (SEPT7 siRNA) of SEPT7. Data represents the mean ± SEM from n = 1,578 (control siRNA) and n = 1,445 (SEPT7 siRNA) bacteria distributed in 4 independent experiments. ns, p > 0.05 by two-tailed Student′s t-test.

**Supplementary Figure 9 related to Figure 4.**
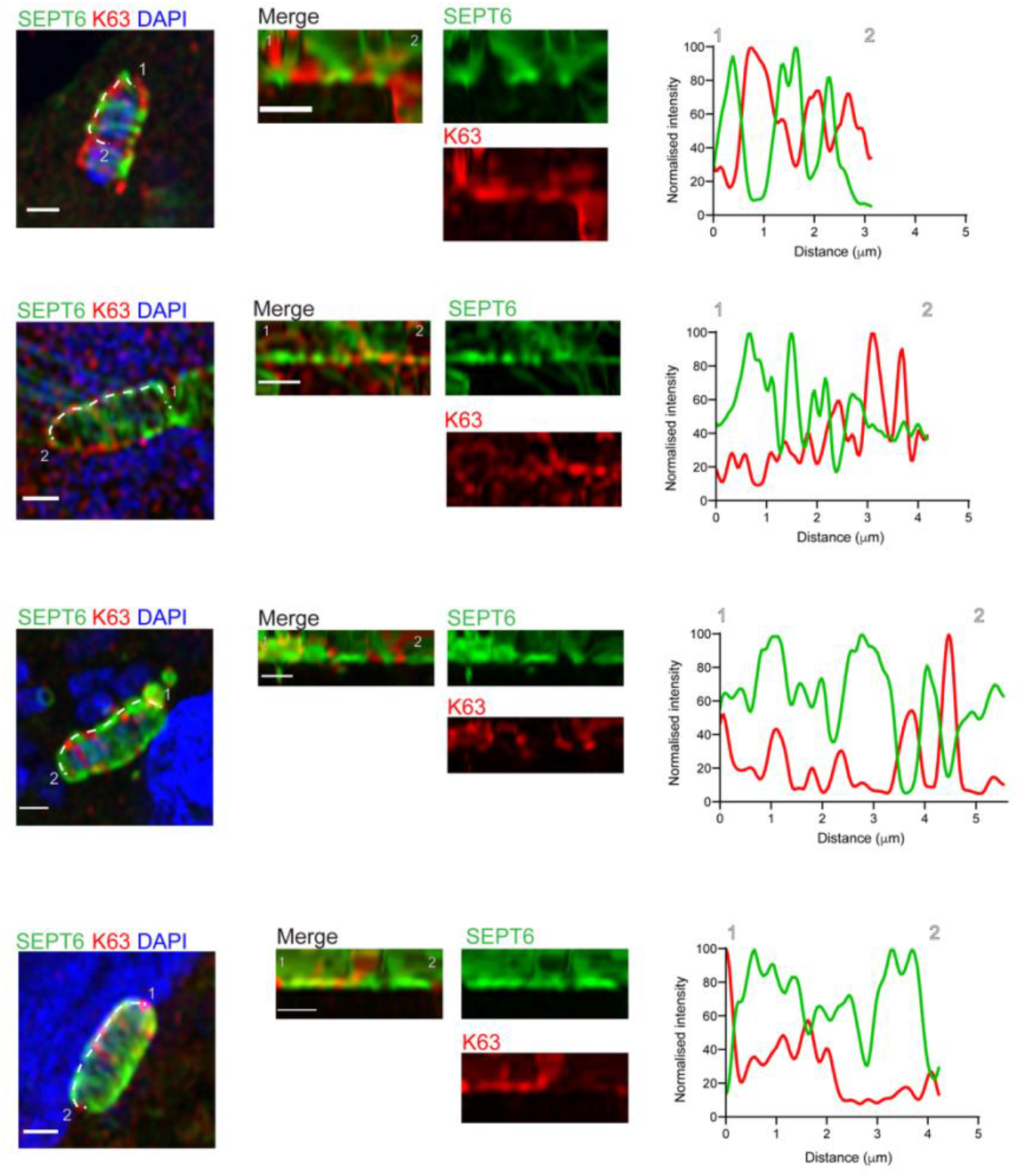
Septins and K63 polyubiquitin form separate microdomains on the cell surface of *S. flexneri*. Airyscan confocal images showing the formation of separate microdomains of GFP-SEPT6 and K63 polyubiquitin on the surface of *S. flexneri* (left panels). Central panels represent the GFP-SEPT6 and K63 polyubiquitin microdomains of the septin cages from left panels. 1 and 2 mark the beginning and end of the dashed white line from left panels. On the right panels it is represented the fluorescence intensity profiles of GFP-SEPT6 and K63 polyubiquitin across the line from central panels. Scale bar, 1 μm.

**Supplementary Table 1.**
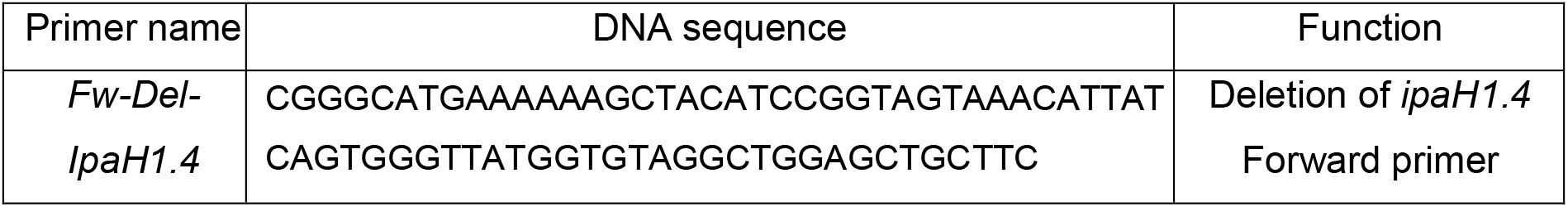
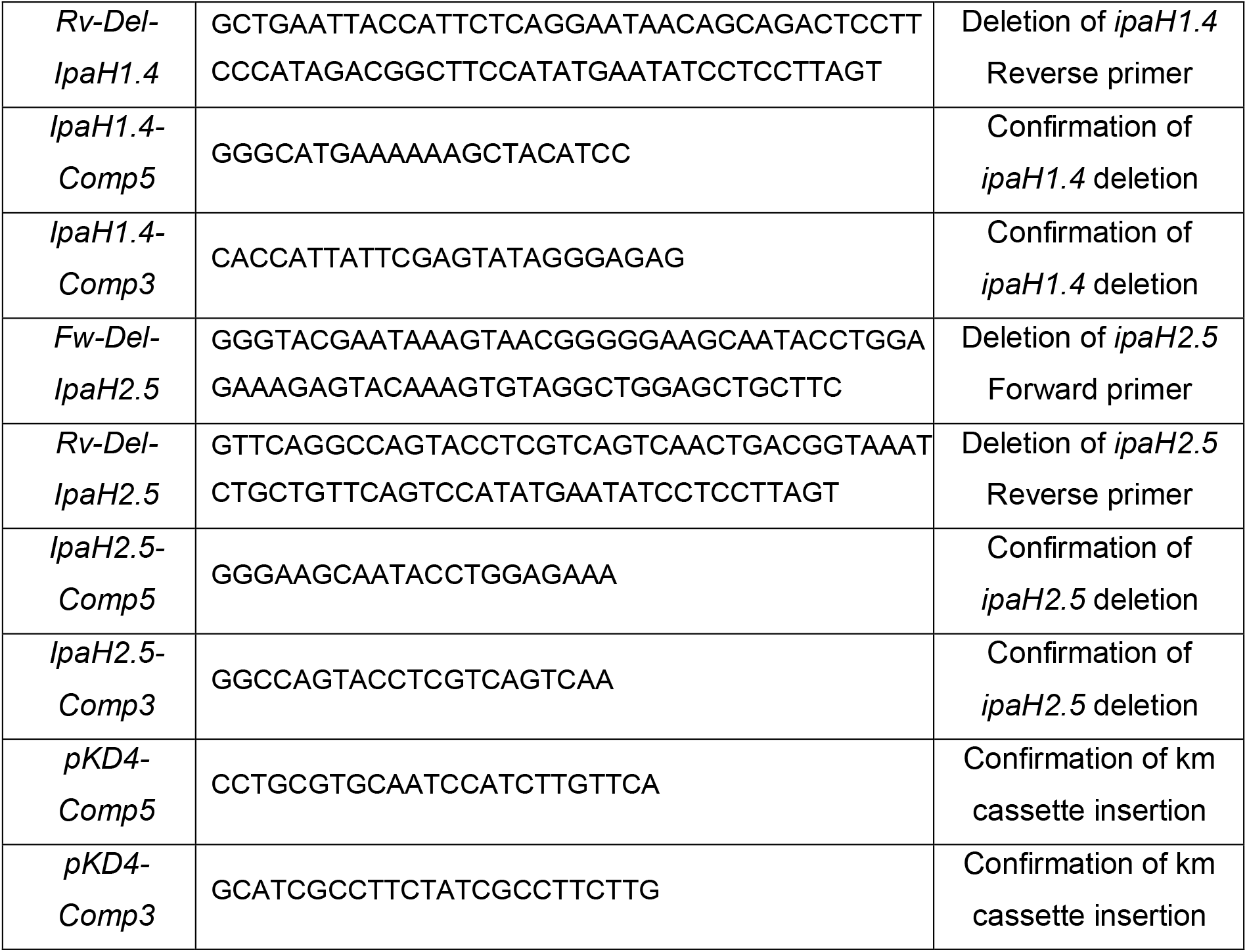
Oligonucleotides used in this study.

## MOVIE LEGENDS

**Movie S1**. Correlative light and cryo-SXT movie related to Figure 2B.

**Movie S2**. Correlative light and cryo-SXT movie related to Figure 3B.

**Movie S3**. Time-lapse epifluorescence movie related to Fig. S5A. Images were acquired as 11 z-stacks every 4 min. Images were deconvoluted using ZEN Blue software and max projected. Scale bar, 5 µm.

**Movie S4**. Time-lapse epifluorescence movie related to Fig. S5B. Images were acquired as 10 z-stacks every 3 min. Images were deconvoluted using ZEN Blue software and max projected. Scale bar, 5 µm.

**Movie S5**. Time-lapse Airyscan confocal movie related to Fig. S6A. Images were acquired using Airyscan fast mode as 12 z-stacks every 5 min. Images were processed using “3D Airyscan processing” using ZEN Black software and max projected. White arrow indicates the interaction between GFP-SEPT6 and mCherry-LC3B mediating the recruitment of mCherry-LC3B to the *Shigella* septin cage. Scale bar, 1 µm.

**Movie S6**. Time-lapse Airyscan confocal movie related to Fig. S6B. Images were acquired using Airyscan fast mode as 10 z-stacks every 4 min. Images were processed using “3D Airyscan processing” using ZEN Black software and max projected. Scale bar, 1 µm.

## Notes

### Competing Interest Statement

The authors have declared no competing interest.

