## Supplementary material for "Septins and K63 chains form separate bacterial microdomains during autophagy of entrapped *Shigella*": ImageJ macros used

**ImageJ macro 1**

//Macro Flatten_bacterial_contours

//This macro is used to linearise the bacterial contour on a MAX projection with 4 channels
//IT should be used after the macro draw_elipse. The ROI set used should contain the two intersection
//points identified with the macro draw_elipse and then the reduced contour also obtained with the macro draw_elipse.

//Writen by Ana T. Lopez-Jimenez at Serge Mostowy lab (@ LSHTM) in 2021.

```
name = getTitle();
nameminustif = substring(name, 0, indexOf(name, ".tif"))
nameduplicate = nameminustif + "-1.tif"
dir = getDirectory("image");

run("Split Channels");


run("Line Width...", "line=0.5");
selectWindow("C1-" + name);
run("Duplicate...", " ");


roiManager("Select", 2);
run("Area to Line", "title=["+"C1-"+ nameduplicate +"]" );
run("Straighten...", "title=["+"C1-"+ nameduplicate +"] line=50");
saveAs("Tiff", dir + File.separator + "C1-" + "_straight");


selectWindow("C1-" + name);
run("RGB Color");
setColor("white");
roiManager("Select", 0);

run("Draw", "slice");
setColor("yellow");
roiManager("Select", 1);

run("Draw", "slice");
roiManager("Select", 2);
run("Area to Line", "title=["+"C1-"+ name +"]" );
run("Straighten...", "title=["+"C1-"+ name +"] line=50");
saveAs("Tiff", dir + File.separator + "C1-" + "_straight_dots");

close();

run("Line Width...", "line=0.5");
selectWindow("C2-" + name);
```

```
run("Duplicate...", " ");


roiManager("Select", 2);
run("Area to Line", "title=["+"C1-"+ nameduplicate +"]" );
run("Straighten...", "title=["+"C1-"+ nameduplicate +"] line=50");
saveAs("Tiff", dir + File.separator + "C2-" + "_straight");

selectWindow("C2-" + name);
run("RGB Color");
setColor("white");
roiManager("Select", 0);
run("Draw", "slice");
setColor("yellow");
roiManager("Select", 1);
run("Draw", "slice");
roiManager("Select", 2);
run("Area to Line", "title=["+"C2-"+ name +"]" );
run("Straighten...", "title=["+"C2-"+ name +"] line=50");
saveAs("Tiff", dir + File.separator + "C2-" + "_straight_dots");
close();


run("Line Width...", "line=0.5");
selectWindow("C3-" + name);
run("Duplicate...", " ");


roiManager("Select", 2);
run("Area to Line", "title=["+"C3-"+ nameduplicate +"]" );
run("Straighten...", "title=["+"C3-"+ nameduplicate +"] line=50");
saveAs("Tiff", dir + File.separator + "C3-" + "_straight");

selectWindow("C3-" + name);
run("RGB Color");
setColor("white");
roiManager("Select", 0);
run("Draw", "slice");
setColor("yellow");
roiManager("Select", 1);
run("Draw", "slice");
roiManager("Select", 2);
run("Area to Line", "title=["+"C3-"+ name +"]" );
run("Straighten...", "title=["+"C3-"+ name +"] line=50");
saveAs("Tiff", dir + File.separator + "C3-" + "_straight_dots");
close();

run("Line Width...", "line=0.5");
selectWindow("C4-" + name);
run("Duplicate...", " ");


roiManager("Select", 2);
run("Area to Line", "title=["+"C1-"+ nameduplicate +"]" );
run("Straighten...", "title=["+"C1-"+ nameduplicate +"] line=50");
```

```
saveAs("Tiff", dir + File.separator + "C4-" + "_straight");

selectWindow("C4-" + name);
run("RGB Color");
setColor("white");
roiManager("Select", 0);
run("Draw", "slice");
setColor("yellow");
roiManager("Select", 1);
run("Draw", "slice");
roiManager("Select", 2);
run("Area to Line", "title=["+"C4-"+ name +"]" );
run("Straighten...", "title=["+"C4-"+ name +"] line=50");
saveAs("Tiff", dir + File.separator + "C4-" + "_straight_dots");

close();
```

**ImageJ macro 2**

```
//Macro draw_elipse

//This macro is used to analyse the recruitment of host factors to a bacterial surface.
//It fits a binary mask (corresponding to the bacteria plus host factors to analyse) to an
elipse.
//Then,it finds the intersection of the max axis of the elipse and the reduced contour of the
mask
//and saves it as ROI sets.

//Writen by Ana T. Lopez-Jimenez at Serge Mostowy lab (@ LSHTM) in 2021.

name = getTitle();
dir = getDirectory("image");


run("Set Measurements...", " area mean centroid fit decimal=6");
  roiManager("Select", 0);
  run("Enlarge...", "enlarge=-0.10");
  roiManager("Update");
  run("Set Scale...", "distance=1 known=1 pixel=1 unit=pixel");
  run("Fit Ellipse");
  roiManager("Measure");
  run("RGB Color");


  for(i=0; i<nResults; i++) {
      x=getResult('X',i);
      y=getResult('Y',i);
      d=getResult('Major',i);
      a = getResult('Angle',i)*PI/180;
      setColor("red");
      makeLine(x+(d/2)*cos(a),y-(d/2)*sin(a),x-(d/2)*cos(a),y+(d/2)*sin(a));
      d=getResult('Minor',i);
      a=a+PI/2;
      roiManager("Add");

  }
roiManager("Save", dir + File.separator + "_ROI_reduced_object_line.zip");

run("Select None");
run("Duplicate...", " ");
setBackgroundColor(0, 0, 0);
run("Select All");
run("Clear", "slice");
roiManager("Select", 0);
run("Draw", "slice");
run("Select All");
saveAs("Tiff", dir + File.separator + "1.tif");


run("Select None");
run("Duplicate...", " ");
setBackgroundColor(0, 0, 0);
```

```
run("Select All");
run("Clear", "slice");
roiManager("Select", 1);
run("Draw", "slice");
run("Select All");
saveAs("Tiff", dir + File.separator + "2.tif");

imageCalculator("AND create", "1.tif", "2.tif");
run("Make Binary");
saveAs("Tiff", dir + File.separator + "3.tif");
run("Set Measurements...", "area mean centroid fit display redirect=None decimal=3");
run("Analyze Particles...", "display exclude clear add");
roiManager("Deselect");
roiManager("Save", dir + File.separator + "_ROI_intersection.zip");
```
